# Cellular proteomic profiling using proximity labelling by TurboID-NES in microglial and neuronal cell lines

**DOI:** 10.1101/2022.09.27.509765

**Authors:** Sydney Sunna, Christine Bowen, Hollis Zeng, Sruti Rayaprolu, Prateek Kumar, Pritha Bagchi, Qi Guo, Duc M. Duong, Sara Bitarafan, Aditya Natu, Levi Wood, Nicholas T. Seyfried, Srikant Rangaraju

## Abstract

Different brain cell types play distinct roles in brain development and disease. Molecular characterization of cell-specific mechanisms using cell type-specific approaches at the protein (proteomic) level, can provide biological and therapeutic insights. To overcome the barriers of conventional isolation-based methods for cell type-specific proteomics, *in vivo* proteomic labeling with proximity dependent biotinylation of cytosolic proteins using biotin ligase TurboID, coupled with mass spectrometry (MS) of labeled proteins, has emerged as a powerful strategy for cell type-specific proteomics in the native state of cells without need for cellular isolation. To complement *in vivo* proximity labeling approaches, *in vitro* studies are needed to ensure that cellular proteomes using the TurboID approach are representative of the whole cell proteome, and capture cellular responses to stimuli without disruption of cellular processes. To address this, we generated murine neuroblastoma (N2A) and microglial (BV2) lines stably expressing cytosolic TurboID to biotinylate the cellular proteome for downstream purification and analysis using MS. TurboID-mediated biotinylation captured 59% of BV2 and 65% of N2A proteomes under homeostatic conditions. TurboID expression and biotinylation minimally impacted homeostatic cellular proteomes of BV2 and N2A cells, and did not affect cytokine production or mitochondrial respiration in BV2 cells under resting or lipopolysaccharide (LPS)-stimulated conditions. These included endo-lysosome, translation, vesicle and signaling proteins in BV2 microglia, and synaptic, neuron projection and microtubule proteins in N2A neurons. The effect of LPS treatment on the microglial proteome was captured by MS analysis of biotinylated proteins (>500 differentially-abundant proteins) including increased canonical pro-inflammatory proteins (Il1a, Irg1, Oasl1) and decrease anti-inflammatory proteins (Arg1, Mgl2).

## 2.0 INTRODUCTION

The brain is a complex organ possessing heterogeneous populations of neurons, glia and vascular cells. The orchestration of interactions within cell types (cell autonomous) and between cell types (non-cell autonomous) support higher-level processes critical to development, psychiatric conditions, and neurodegeneration. Protein-level analyses using mass spectrometry (MS) expands upon other systems-level analyses, including genomics and transcriptomics, in its ability to profile total protein abundances, along with post-translational modifications, and resolve protein-level changes occurring in subcellular compartments. A central challenge to neuroproteomics is the difficulty in obtaining cell-type specific proteomes from brain tissue.

Traditional approaches to isolating cell-type specific proteomes for MS including fluorescence activated cell sorting (FACS) and magnetic activated cell sorting (MACS) require fresh brain tissue, and the harsh and laborious processing itself poses challenges.^1,2^ Majority of adult neurons do not survive the isolation process, and sampling bias for healthier non-neuronal brain cells able to withstand the isolation process limits proteomic profiling in disease states. Additionally, contamination from proteins derived from other cell types persists. The challenges to maintaining cellular integrity with isolation methods motivated the field to innovate novel methods of applying cell-type specific labeling to *in-vitro* and *in-vivo* systems.

One emergent approach to achieve cell type-specific proteomic labeling uses BioOrthogonal Non-Canonical Amino Acid Tagging (BONCAT) in which a mutated methionyl-tRNA synthetase (MetRS) incorporates azidonorleucine (ANL), a methionine analogue, into newly synthesized peptides.^3–5^ Subsequently, cell lysates or brain homogenates undergo click chemistry with biotin-alkyne to biotinylate the ANL-containing peptides. By driving MetRS expression under a cell-specific promoter and enriching biotinylated peptides by streptavidin affinity-purification, BONCAT can purify cell-type specific newly-translated proteins.^6,7^ One advantage of this strategy lies in its ability to label and purify low-abundant and newly-synthesized proteins. A limitation may be low proteomic depth and biases towards proteins with high turnover. The BONCAT approach has thus far been applied to characterize excitatory and inhibitory neurons of mice and rats in both *in-vivo, ex-vivo,* and *in-vitro* contexts.^6–9^ To date, extension to other neuronal and glial cell types has not yet been published.

In contrast to the BONCAT approach which labels only newly synthesized proteins, proximity-labeling techniques rely on biotin ligases which biotinylate nearby interactors promiscuously. BioID is a promiscuous biotin-ligase engineered from the site-specific biotin ligase, BirA, endogenously produced by *Escherichia coli*.^10^ Because BioID is non-toxic, this technology opened up opportunities for *in-vivo* applications, though the reaction kinetics of biotin-labeling takes place over 18-24 hours.^11^ Alice Ting’s group used yeast surface display mediated directed evolution to improve the reaction kinetics of BioID by introducing 15 mutations in the catalytic domain relative to wild-type BirA.^12^ This biotin ligase, termed TurboID, can robustly and promiscuously biotinylate proteomes in living cells and animal models without cellular toxicity, in as little as 10 minutes in cell culture systems.^12,13^ Versatile in its applications, TurboID has been fused to proteins of interest to map protein interactomes, targeted to a subcellular compartments of interest, and has been exported out of the nucleus to label cytosolic proteins.^14–19^ Additionally, to biotin-label proteins proximal to intermembrane contact sites, split-TurboID was recently developed in which two inactive fragments of TurboID reconstitute in the presence of rapamycin.^13,20^ When driven under a cell-type specific promoter, both TurboID and split-TurboID can label cellularly distinct proteomes in their native state for downstream affinity capture and MS. These advancements, which enable robust, targeted, and non-toxic biotinylation of cell-type specific proteomes has yielded promising applications to neuroproteomics. Recently, split-TurboID has been applied to perisynaptic-cleft proteome discovery between astrocyte and neurons *in vivo*.^18,21^ A novel transgenic mouse model for conditional expression of TurboID with a nuclear export sequence (TurboID-NES) was also recently developed to resolve region-specific proteomic signatures of CamKIIa neurons and Aldh1l1 astrocytes in adult mouse brain. ^21,22^ These recent advances position cell type-specific *in-vivo* biotinylation of proteins (CIBOP) as a promising approach to resolve distinct cellular proteomes in different *in vivo* homeostatic and pathological contexts.

In anticipation of *in vivo* proximity labeling applications of TurboID-NES for cell type-specific proteomics, it is important to establish the effects of global cytosolic biotinylation on molecular and cellular processes in mammalian cells. It is also critical to characterize the breadth of cytosolic proteins labeled by TurboID-NES, determine how reflective these proteins are of un-transduced whole cell proteomes and identify any inherent biases of the TurboID-NES approach. In order to support the use of TurboID-NES to label cytosolic proteins which are also relevant to cellular identity, it is important to test if TurboID- mediated proteomics can differentiate two distinct cell types, such as neurons and microglia. Finally, we need to ascertain whether proteomic changes induced by activation stimuli can be efficiently and reliably captured by the TurboID-NES approach. These data are critical for interpreting proteomic results from TurboID-NES proximity labeling studies that aim to label the cellular proteome in mammalian systems *in vitro* and *in vivo*. To answer these questions under controlled experimental conditions, we generated neuroblastoma (N2A) and immortalized microglial (BV2) cell lines that stably express V5- TurboID-NES to label the cellular proteome excluding the nucleus. We examined the extent and coverage of cytosolic proteomic labeling by TurboID-NES in N2A and BV2 cells, under resting and lipopolysaccharide (LPS)-stimulated inflammatory conditions, using label-free quantitation (LFQ) mass spectrometry. We found that TurboID-NES expression preserves ability to resolve distinct cellular proteomes at the MS level under homeostatic states and inflammatory challenge.

## 3.0 RESULTS

### Generation and validation of stably-transduced microglial and neuronal TurboID- NES cell lines

We created a lentiviral vector incorporating the V5-TurboID-NES sequence (Addgene plasmid #107169) including a Green Fluorescence Protein (GFP) separated by a T2A linker, and a puromycin resistance gene (**Fig 1A**). The NES sequence was incorporated to limit biotinylation to the extra-nuclear compartment. We then generated lentiviruses carrying this vector and transduced BV2 mouse microglia and N2A mouse neuroblastoma cells (MOI 5:1) followed by positive selection with puromycin for at least 2 passages and then fluorescent activated cell sorting (FACS) of GFP-positive cells. These sorted cells were then maintained in puromycin for passages to maintain stably-transduced BV2-Turbo and N2A-Turbo lines (**Fig 1B**) but cultured in puromycin-free medium prior to experimentation. After genetically screening successfully transduced cell lines with puromycin, we used western blotting (WB) and immunofluorescence (IF) to confirm robust biotinylation of proteins in cell lysates (**Fig 1C**) and confirm cytosolic localization of V5-TurboID-NES and biotinylated proteins (**Fig 1D, 1E**). WB probing for biotinylated proteins consistently identified few proteins in untransduced controls which likely represent endogenously biotinylated carboxylases such as pyruvate carboxylase (∼130 kDa), 3-methylcrotonyl coA carboxylase (∼75 kDA) and propionyl coA carboxylase (72 kDa).^23^ The results shown in **Fig 1D** and **1E** confirmed the functionality of the NES, and the predominant biotinylation of cytosolic proteins in both BV2-TurboID-NES and N2A- TurboID-NES cells.

**Figure 1.**
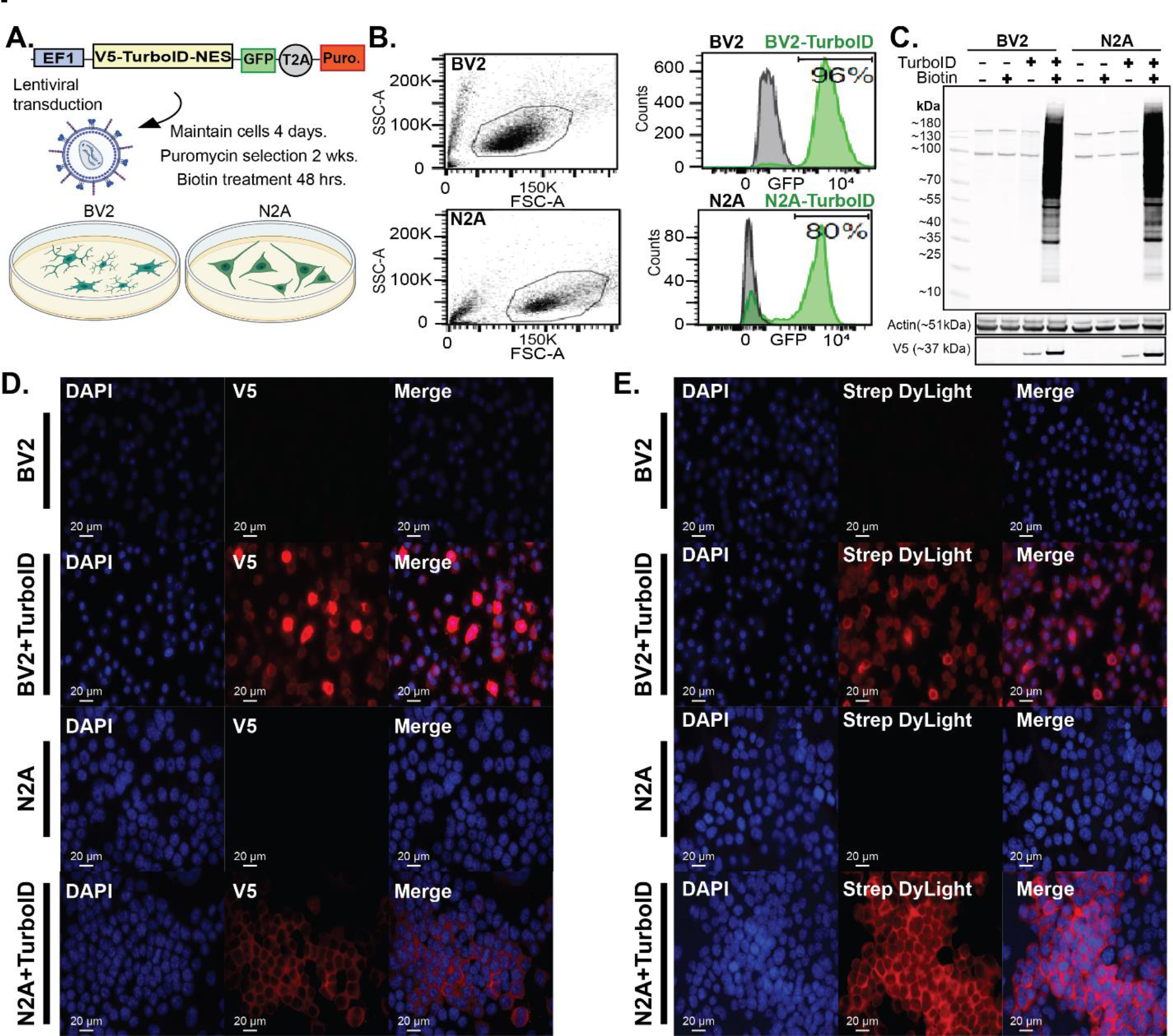
Creation of stably transduced BV2 and N2A cells. **A.** Schematic of transduction. The genetic construct packaged into a lentivirus contains V5-tagged TurboID- NES driven under an EF1 promoter and a GFP sequence separated by a T2A linker. N2A and BV2 cells were transduced and maintained for 4 days prior to 2 weeks of puromycin selection and biotin supplementation in media. **B.** Following puromycin selection, FLOW cytometry confirms GFP positivity in a majority of BV2 (96%) and N2A (80%) cells. **C.** Western blot (WB) of transduced and untransduced cell lysates confirming presence of TurboID (V5) and biotin-dependent robust biotinylation of proteins. Actin was used as a loading control. **D.** Immunofluorescence (IF) confirming cytoplasmic localization of V5- TurboID-NES in transduced cells. **E.** IF confirming cytosolic biotinylation of proteins in transduced BV2 and N2A cells.

### TurboID-NES-based MS captures representative proteomes in mammalian microglial and neuronal cell lines

After confirming stable expression, cytosolic localization and functionality of V5- TurboID-NES in BV2 and N2A cell lines, we prepared cell lysates for LFQ-MS studies. Transduced and untransduced BV2 and N2A cells received 1 µg/mL of LPS or equal volume of phosphate-buffered saline (PBS) in addition to 200 µM biotin supplementation for 48 hours. LPS was used as a general immune stimulus to activate BV2 cells while anticipating marginal or no effects of LPS on N2A. Whole-cell (WC) lysates were prepared in parallel with lysates undergoing streptavidin affinity-purification (AP) to enrich for biotinylated proteins (n = 4 / group) (**Fig 2A**). WC lysates underwent Coomassie staining and were probed with Streptavidin-680 to confirm robust biotinylation of proteins in transduced cell-lines (**Fig 2B, 2C**). Proteins bound to streptavidin beads were boiled and resolved with gel electrophoresis (**Fig 2D, 2E**). Proteins released from streptavidin-beads were probed for biotinylation using streptavidin 680, and silver-stained gels were run to confirm minimal binding of non-biotinylated proteins in untransduced lysates. In BV2 AP lysates, LPS treatment induced differential banding patterns visible in smaller molecular weight proteins (10-40 kDa) (**Fig 2D**). The silver stain and western blots provided evidence for not only the capacity of TurboID-NES to biotinylate proteins altered by LPS treatment, but also the ability to affinity-purify LPS-altered proteins biotinylated by TurboID-NES. Notably, LPS did not alter protein banding patterns in either the silver stain or western blots of N2A samples. After confirming the enrichment of biotinylated proteins in TurboID- NES-transduced BV2 and N2A cell lines, we submitted these same samples for LFQ-MS. Raw LFQ and intensity values are found in **Supplemental Datasheet 1A** (**SD 1A**).

**Figure 2.**
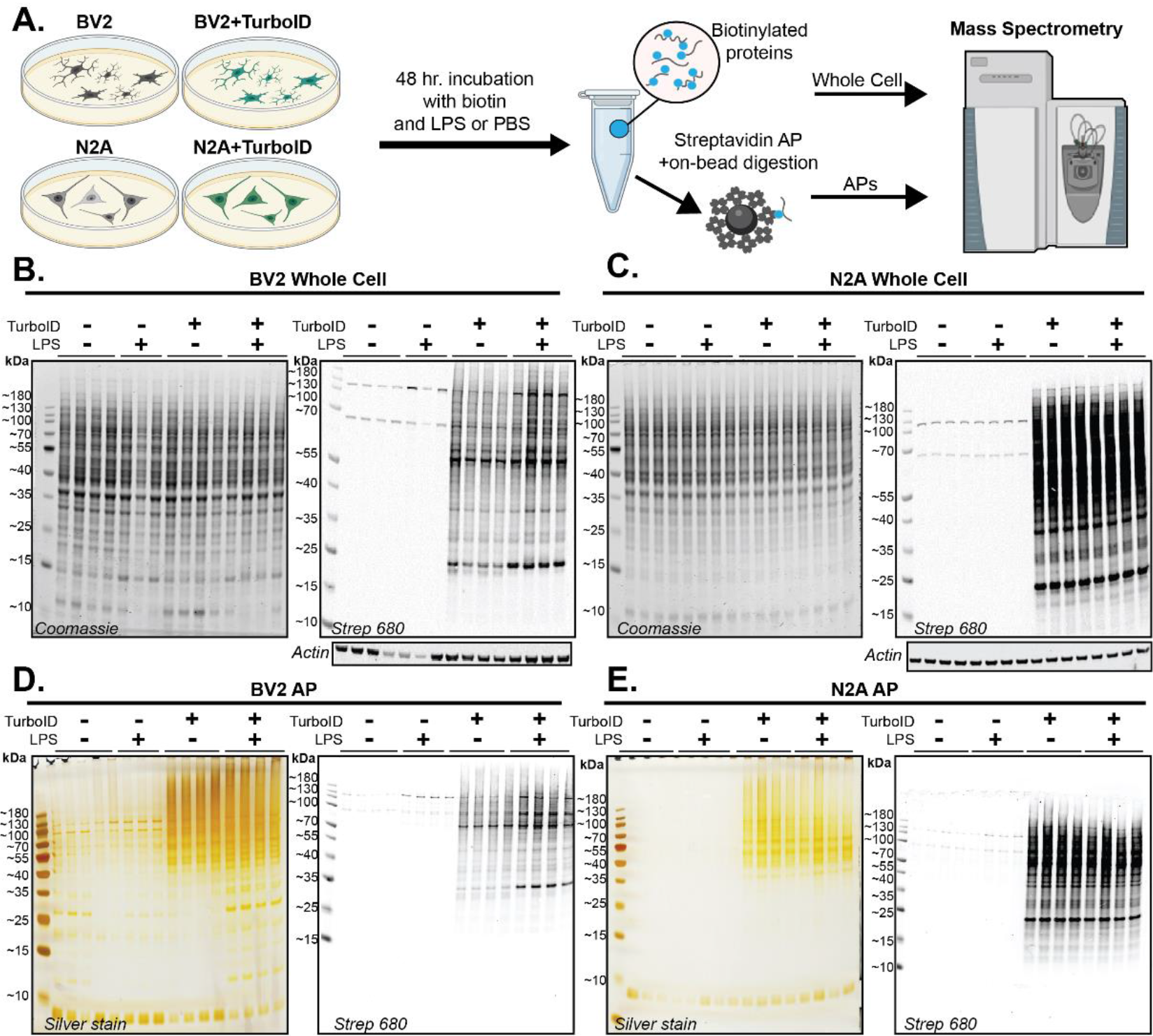
Experimental design and quality control of whole-cell and AP samples prior to MS. **A.** Schematic of experimental design. Transduced and untransduced BV2 and N2A cells were treated with biotin and LPS or PBS for 48 hrs. Whole-cell (WC) lysates and streptavidin- affinity purified (AP) samples were processed in parallel with Mass Spectrometry (MS) **B.** Western blot (WB) and Coomassie confirming biotinylation of proteins in transduced BV2 WC lysates. **C.** WB and Coomassie confirming biotinylation of proteins in transduced N2A WC lysates **D.** WB and silver stain confirming biotinylation of proteins bound to streptavidin beads and specificity of streptavidin beads for biotinylated species in BV2 AP preparations. LPS impacts banding patterns (10-40kDa) in transduced BV2 biotinylated proteins visualized with both Silver Stain and WB. **E.** WB and silver stain confirming biotinylation of proteins bound to streptavidin beads and specificity of streptavidin beads for biotinylated species in N2A AP preparations.

First, we used PCA for dimension reduction of LFQ proteomic data obtained from all whole cell samples and transduced AP samples (**SF 1A, 1B**). LFQ-MS data used for WC- level comparisons are available in **(SD 2A)** and TurboID-normalized intensity values used for AP-level comparisons are available (**SD 2B**). WC samples included BV2 (n = 4), BV2+LPS (n = 3), BV2+TurboID-NES (n = 4), BV2+TurboID-NES+LPS (n = 4) N2A (n = 4), N2A+LPS (n = 4), N2A+TurboID-NES (n = 4), and N2A+LPS+TurboID-NES (n = 4) (**SF 1A**). Due to the high number of missingness in the non-transduced +AP samples that only captures endogenously biotinylated proteins, we performed PCA on +TurboID-NES or +LPS+TurboID-NES BV2 and N2A AP samples (n = 4/group) (**SF 1B**). In the PCA of WC samples, PC1 described ∼32% of variance across WC proteomes and captured cell type differences between BV2 and N2A cells without contribution from TurboID-NES status or LPS treatment (**SF 1A**). The full WC principal component matrix can be found in (**SD 1B**). In PCA of AP samples, PC1 accounted for 48% of the variance and also captured BV2 N2A cell type differences. PC2 accounted for 10% variance and captured the effect of LPS only in BV2 cells and not in N2A cells (**SF 1B**). The AP principal component matrix can be found in (**SD 1C**). These PCA analyses of WC as well as AP proteomes confirmed that the cell type differences between BV2 and N2A proteomes, rather than TurboID-NES expression or proteomic biotinylation, explained the majority of variance in our data. Importantly, biotin-enriched proteomes from AP samples successfully resolved cell type differences as well as LPS-treatment effects within the BV2 cells.

To assess the global proteomic differences and similarities between untransduced WC proteomes (BV2 or N2A whole cell, n =4 / group) and their TurboID-NES-transduced and biotin-enriched counterparts (BV2+TurboID-NES_AP_ or N2A+TurboID-NES_AP_, n = 4 / group), we performed k-means clustering analysis (**Fig 3A**). We identified 6 distinct clusters using the elbow method for cluster number optimization (**SF 2**). We identified clusters of proteins preferentially abundant in WC proteomes (Cluster 1), and clusters of proteins abundant in TurboID-NES AP samples and their WC counterparts (Clusters 3 and 6). Interestingly, we also identified clusters of proteins distinct to TurboID-NES AP samples which TurboID-NES-mediated biotinylation showed preferential abundance for (Clusters 2 and 4). Finally, we identified a cluster of proteins which are shared in enrichment between TurboID-NES AP cells (Cluster 5). LFQ-MS identified 3,064 proteins in BV2 proteomes, of which TurboID-NES biotinylated 1,815 or ∼59% LFQ MS identified 3,173 proteins in N2A proteomes, of which TurboID-NES biotinylated a total of 2,056 proteins or ∼65% (**Fig 3B**).

**Figure 3.**
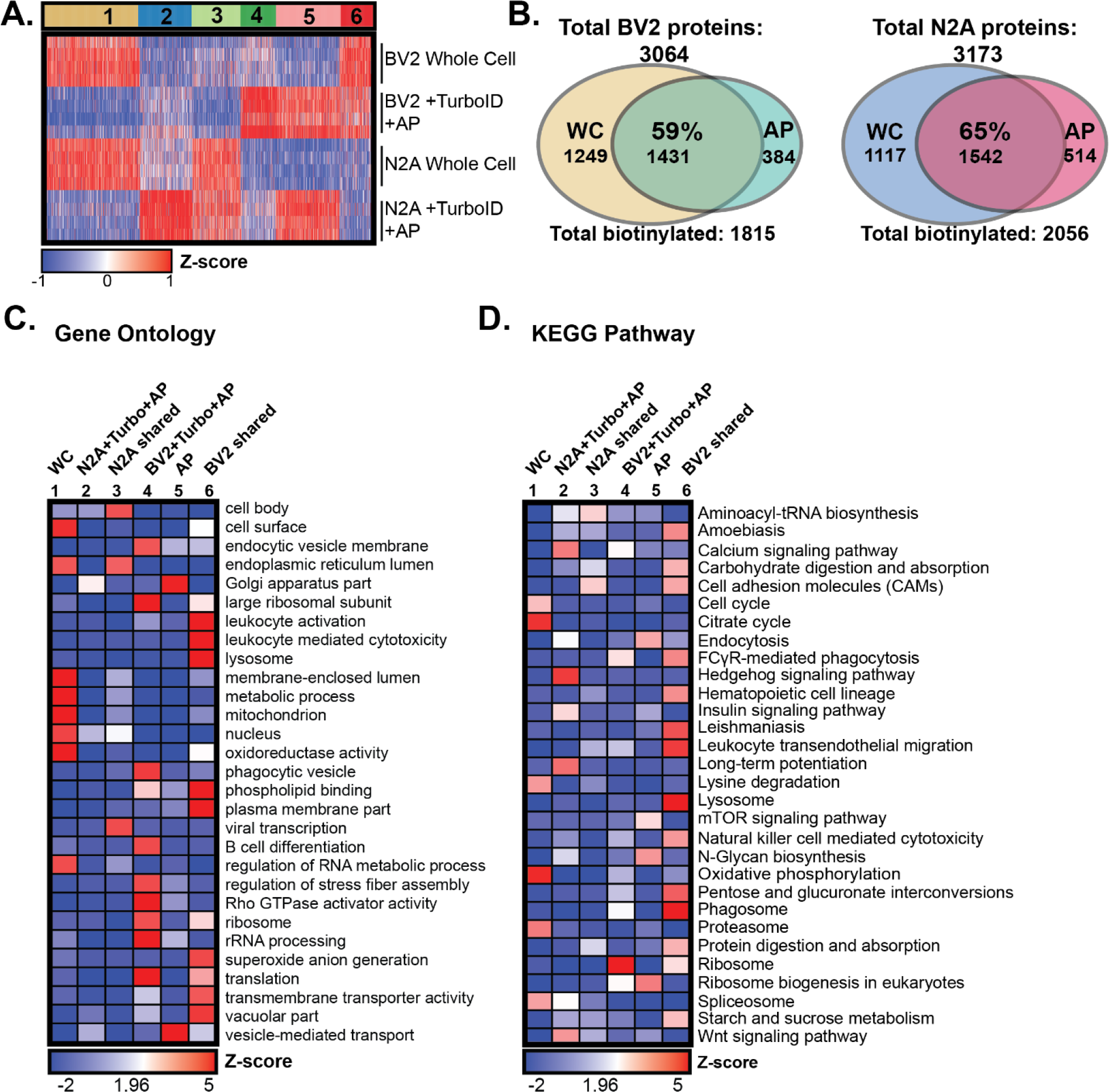
Global profiling of TurboID labeling in microglial and neuronal cell lines. A. K-means clustered heatmap representation of LFQ intensity data of proteins identified by MS in WC and biotinylated and AP proteomes (n = 4 per experimental group) in microglial and neuronal cell lines. 6 distinct clusters represent labeling profile of TurboID, including proteomes enriched in whole-cell preparations, AP preparations, and by cell type. B. Venn diagram of protein counts identified in BV2 WC samples and BV2+TurboID+AP samples and N2A WC and N2A+TurboID+AP samples. In BV2 cells, TurboID-NES labels ∼59% of the total proteins captured by LFQ MS. In N2A cells, TurboID-NES captures ∼65% of the total proteins identified by LFQ MS. **C.** Functional annotation of gene-set enrichment based on gene-ontology (GO) over-representation analysis (ORA). Heatmap color intensity is based on Z-score enrichment across the 6 clusters. **D.** Heatmap representation and functional annotation of clusters derived from the KEGG pathways.

We performed gene set enrichment analyses (GSEA) using each of the 6 cluster- sets compared with the background proteome of all proteins identified in the LFQ-MS dataset (∼2187 proteins; **SD 2A**), using KEGG and Gene-Ontology (GO) reference databases.^24–28^ GSEA of the WC Cluster 1 (630 proteins) showed enrichment of nuclear, mitochondrial, cell surface and RNA metabolic proteins (**Fig 3C, D**). Cluster 1 represented proteins derived from compartments less accessible to biotinylation by TurboID-NES in both cell types. In contrast, Cluster 5 (425 proteins) contained proteins selectively abundant in both BV2 and N2A transduced AP samples. Cluster 5 represented a group of proteins preferentially labeled by TurboID-NES in both cell types. GSEA of Cluster 5 showed enrichment of Golgi apparatus, vesicle-mediated transport and endocytosis related proteins, indicating that TurboID-NES preferentially biotinylated Golgi, endocytic, secretory compartments in both cell types. As TurboID-NES trafficked throughout the cytosol, it came in contact with vesicular compartments and labeled proteins involved in vesicular transport. Because vesicle-mediated transport is a biologically conserved function inherent to both microglial and neuronal cells, we expected that the abundance of vesicular-trafficking proteins labeled by TurboID-NES would not differ by cell-type.

In contrast to Clusters 1 and 5 which primarily differentiated WC from AP samples, we identified cell type-specific clusters (Clusters 3 and 6) that were highly abundant in either N2A or BV2 cells, regardless of WC or AP status. Cluster 3 (323 proteins), the ‘N2A shared’ cluster showed a high abundance of proteins shared between WC and transduced AP N2A cells. The ‘N2A shared’ cluster contained proteins enriched in endoplasmic reticulum, cell body and viral transcription proteins, potentially explained by the rapidly-dividing stem-cell-like origin of N2A cells. Cluster 6 (226 proteins), the ‘BV2 shared’ cluster, was highly abundant in WC and AP samples from BV2 cells. The ‘BV2 shared cluster’ was enriched in proteins associated with immune cell (leukocyte activation and trafficking), hematopoietic lineage, lysosomal, plasma membrane, translation, vacuole, metabolism and phagosome functions as would be expected for BV2 microglial cells.

Clusters 2 and 4 showed cell type-specific proteomic differences apparent only in the AP samples but not in the WC lysates. These may be proteins which are expressed at low-levels in the whole cell, and are enriched by TurboID-NES labeling and AP. These clusters therefore represented proteomic features of N2A and BV2 cell types that were readily revealed by the TurboID-NES approach and are less apparent at the level of WC proteomes, potentially via preferential access of TurboID-NES to specific cellular compartments. Cluster 2 (350 proteins) was highly abundant in N2A AP samples compared to BV2 AP samples, and were enriched in proteins involved in long-term potentiation, calcium signaling, Hedgehog signaling, consistent with ontologies expected in neuronal-origin N2A cells. Cluster 4 (233 proteins), on the other hand, was highly abundant in BV2 AP samples and were enriched in endocytosis, ribosomal subunits, phagocytosis, Rho GTPase activity, rRNA processing and translation terms, consistent with a phenotype expected in immune and phagocytic cells such as BV2 microglia. The GSEA of clusters 2 and 4 show that TurboID-NES mediated biotinylation can enrich proteins in N2A and BV2 proteomes, respectively, which are not readily distinguished at the WC level.

Taken together, these proteomic analyses of N2A and BV2 TurboID-NES AP proteomes along with respective WC proteomes, showed that TurboID-NES proteome was representative of the whole cell proteome. In addition, TurboID-NES preferentially biotinylated several classes of proteins shared across mammalian cell types, as well as proteins unique to microglial or neuronal cell type origin. While many of these biotinylated proteins captured changes apparent at the WC level, several cell type-specific proteomic differences were more apparent in the TurboID-NES AP proteomes as compared to the WC proteomes.

### TurboID-NES over-expression has minimal impact on cellular proteomic and functional profiles of BV2 and N2A cells

Lentiviral transduction and over-expression of TurboID-NES in mammalian cells could potentially impact basic cellular functions and phenotypes, and therefore could have confounding implications for *in-vivo* applications of TurboID-NES. To test whether TurboID- NES expression impacts cellular phenotypes under both homeostatic and inflammatory conditions, we compared whole cell transduced and untransduced proteomes, collected culture supernatants for cytokine profiling in response to LPS challenge, and we assessed the respiratory activity of living BV2 cells. To test the hypothesis that TurboID-NES expression itself impacts the WC proteome, we performed differential expression analysis (DEA) comparing BV2+TurboID-NES_WC_ – BV2_WC_ (**Fig 4A**) and N2A+TurboID-NES_WC_ - N2A_WC_ (**Fig 4B**). Only 53 BV2 proteins and 74 N2A proteins, including TurboID-NES, were significantly changed with TurboID-NES expression out of 2,187 total proteins. The DEA comparing transduced and untransduced BV2 or N2A proteomes can be found in (**SD 2C, SD 2D**). This proteomic result provided evidence that TurboID-NES expression minimally impacted WC proteomes of BV2 and N2A cells under resting conditions. The small sizes of the whole cell TurboID-DEP input lists did not yield gene ontology terms in the over- representation analysis. However, the top 5 increased terms from the DEA comparing BV2+TurboID-NES_WC_ – BV2_WC_ included Histone H1.0; H1f0, phosphoserine phosphatase Psph, and kinases; Adrbk1, and Prkab1 and TurboID itself. The 5 most decreased proteins with TurboID transduction in BV2 include vesicle membrane protein, Vat1l, Rho-related GTP binding protein, Rhoc, mitochondrial tRNA ligase Tars2, U3 small nucleolar RNA-associated protein, Utp14a, and cell adhesion molecule 1, Cadm1. When assessing the impact of TurboID expression on N2A WC proteomes, N2A+TurboID-NES_WC_ - N2A_WC_, the DEA revealed the top 5 increased proteins included TurboID, dehydrogenase / reductase family member 7, Dhrs7, Niban, Fam129a, Nuclear valosin, Nvl, and disabled homolog 2, Dab2. The 5 most decreased proteins with TurboID transduction in N2A included putative methyltransferase Nsun7, COBW domain containing protein 1, Cbwd1, putative helicase MOV-10, Mov10, and AHNAK nucleoprotein 2, Ahnak2. Taken together, TurboID transduction in BV2 and N2A cells impacts the abundance of 2% and 3% of the proteins identified in the WC proteome, respectively. Although a minority of proteins are significantly impacted by TurboID, we did observe modest yet significant alterations in nuclear-associated proteins in both cell types, including Histone H1.0 (-Log_10_ p value = 1.34, Log2FC = 2.98) in BV2 and Nuclear valosin-containing protein, Nvl (-Log_10_ p value = 1.33, Log2FC = 1.10) in N2A.

**Figure 4.**
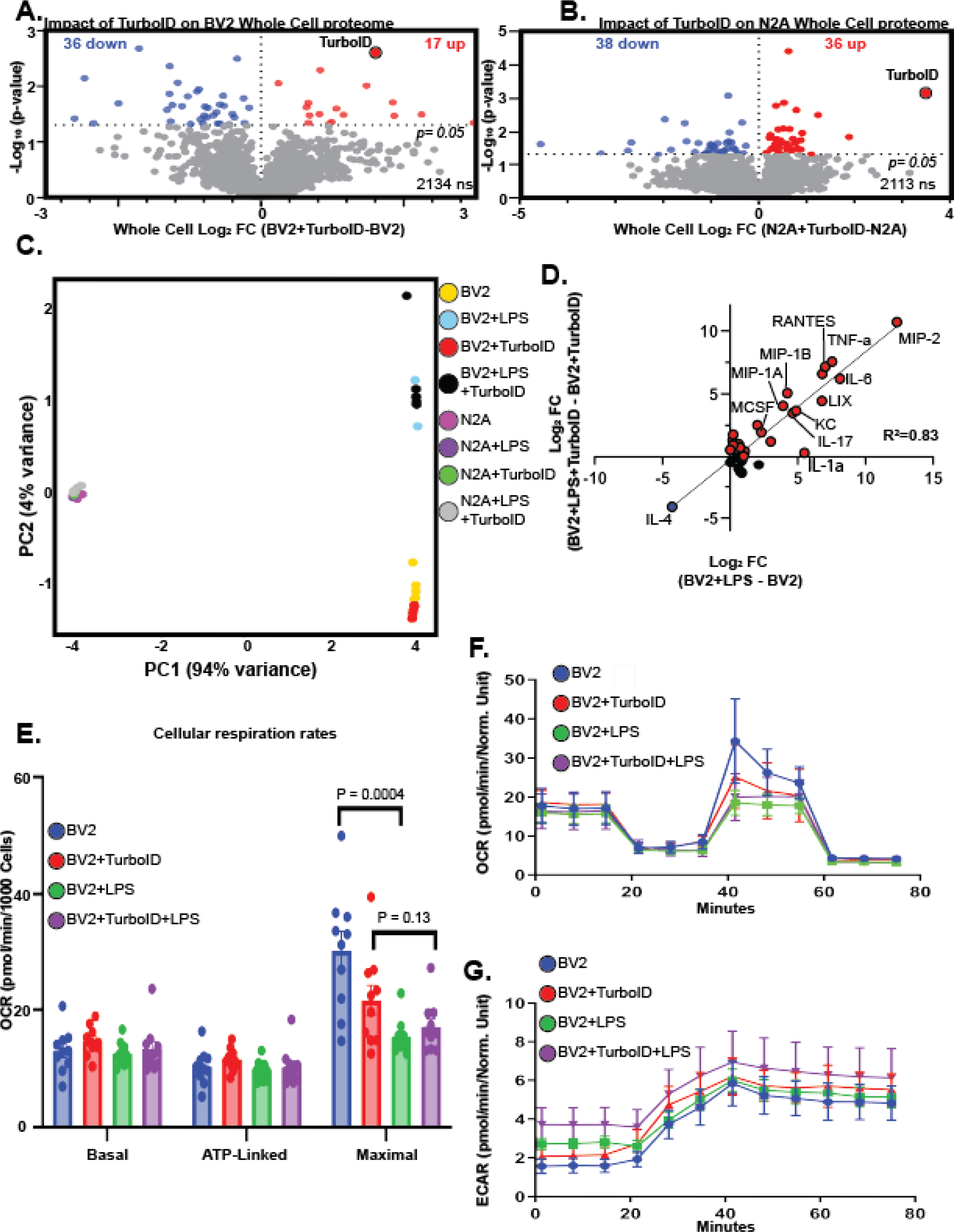
TurboID does not impact cellular phenotypes. **A.** Differential expression analysis (DEA) comparing WC transduced BV2 cell lysates with untransduced BV2 cell lysates identifies 53 Differentially expressed proteins (DEPs) in BV2 with TurboID expression. **B.** DEA comparing WC transduced and untransduced N2A Identifies 74 DEPs **C.** PCA of cytokine profiles derived from cultured supernatants indicates PC1 captures 94% of the variance across samples, accounting for differences in cell type. PC2 captures 4% of the variance, accounting for LPS impact on BV2 cytokines. **D.** Linear regression of LPS cytokine fold-change induction of untransduced (x-axis) and transduced (y-axis) BV2 cells demonstrates high correlation (R^2^=0.83). **E.** Bar graph depicting Basal, ATP-linked and Maximal cellular respiration rates of transduced and untransduced BV2 cells. LPS significantly decreases the maximal respiration of untransduced BV2 cells (p=0.0004) and decreases maximal respiration in transduced BV2 cells, though this finding is statistically insignificant (p=0.13). **F**. Oxygen consumption rate (OCR) traces for transduced and LPS exposed BV2 cells highlight LPS response in maximal respiration. **G**. Extracellular acidification rate (ECAR) highlights the basal glycolytic rate as higher when cells are exposed to LPS but are not impacted by transduction status.

In applying the CIBOP approach to immune cells and inflammatory disease models, it is important to confirm that TurboID expression has a minimal impact on both homeostatic and inflammatory cytokine release profiles. Then, we collected supernatants from TurboID-NES transduced and untransduced BV2 and N2A cells in response to 48 hours of LPS challenge or PBS (n = 6/group) for profiling of 31 cytokines using a Luminex multiplexed immune assay. The raw intensity values, proceeding the subtraction of average background intensity, are found in (**SD 1D**). PCA of secreted cytokines across all samples showed that the cellular identity of cytokines drives a majority of variance (captured by PC1; 94% variance) and that BV2-secreted cytokines robustly responded to LPS while N2A cells did respond to LPS based on secreted cytokine profiles (captured by PC2; 4% variance). The complete principal component matrix can be found in (**SD 1E**).

Importantly, we also observed no separation of BV2 or N2A cells based on TurboID-NES status (**Fig 4C**). To test if TurboID-NES expression impacts the magnitude of cytokine abundance in the presence of LPS, we compared the fold-induction of LPS-driven cytokine abundance changes between transduced and untransduced BV2 cytokines (**Fig 4D**). We observed strong concordance in LPS effects regardless of TurboID-NES status (R^2^=0.83). The secretion of pro-inflammatory cytokines (e.g., macrophage inflammatory protein 2, interleukin 6, and tumor necrosis factor alpha) was increased to comparable extents in response to LPS while anti-inflammatory cytokine (Il-4) was suppressed by LPS to the same extent in BV2 control and BV2 +TurboID-NES cell lines (**Fig 4D**). This result confirmed that TurboID-NES expression did not significantly influence the cytokine release profiles of BV2 cells which received LPS.

To determine if either LPS or TurboID-NES expression impacted cellular respiration rates in transduced or untransduced BV2 cells, we used Seahorse assays of cellular respiration, oxygen consumption rate (OCR), and extracellular acidification rate (ECAR) as a measure of glycolytic activity (**Fig 4E-G**). LPS significantly decreased the maximal respiration of untransduced BV2 cells (p = 0.0004), and we also observed a decrease in the maximal respiration in transduced BV2 cells, though the change was not significant (p = 0.13) (**Fig 4E**). Neither LPS challenge nor transduction status significantly impacted OCR or ECAR in BV2 cells. Taken together, TurboID-NES expression in BV2 and N2A cells had a minimal impact on WC proteomes and does not impact LPS-driven cytokine release. TurboID-NES expression had no significant impact on homeostatic cellular respiration nor glycolytic activity. These *in vitro* findings are of critical importance in interpreting results derived from TurboID-NES-based proteomics by confirming absence of undesired effects of TurboID-NES over-expression in mammalian cells.

### TurboID-NES biotinylates a variety of subcellular compartments within BV2 and N2A cells including several neurodegenerative disease-relevant proteins

The utility of expressing TurboID under cell-type specific promoters *in-vivo* and the consequential purification of cellularly distinct proteins from total brain homogenate lies within the ability of TurboID-NES to label cellularly distinct proteins with disease relevance. Before comparing TurboID-NES-labeled proteomes of BV2 and N2A cells, we first assessed the enrichment of proteins biotinylated by TurboID-NES over endogenously biotinylated proteins. We used DEA to compared AP proteomes from untransduced BV2 lysates with TurboID-NES-transduced BV2 lysates (BV2+TurboID-NES_AP_ – BV2_AP_), which showed that TurboID-NES biotinylated 1754 proteins, whereas 10 endogenously biotinylated proteins appear in the untransduced AP proteome (**Fig 5A; SD 2E**). DEA comparing AP proteomes from untransduced N2A lysates with TurboID-NES-transduced N2A lysates (N2A+TurboID-NES_AP_ – N2A_AP_) showed that TurboID-NES biotinylated 2011 proteins, whereas 39 endogenously biotinylated proteins appeared in the untransduced AP proteome (**Fig 5B; SD 2F**). TurboID-NES expression in BV2 and N2A cell lines, and streptavidin based affinity purification yielded a robustly biotinylated proteome sufficient to over-come the background of endogenously-biotinylated proteins in untransduced AP samples.

**Figure 5.**
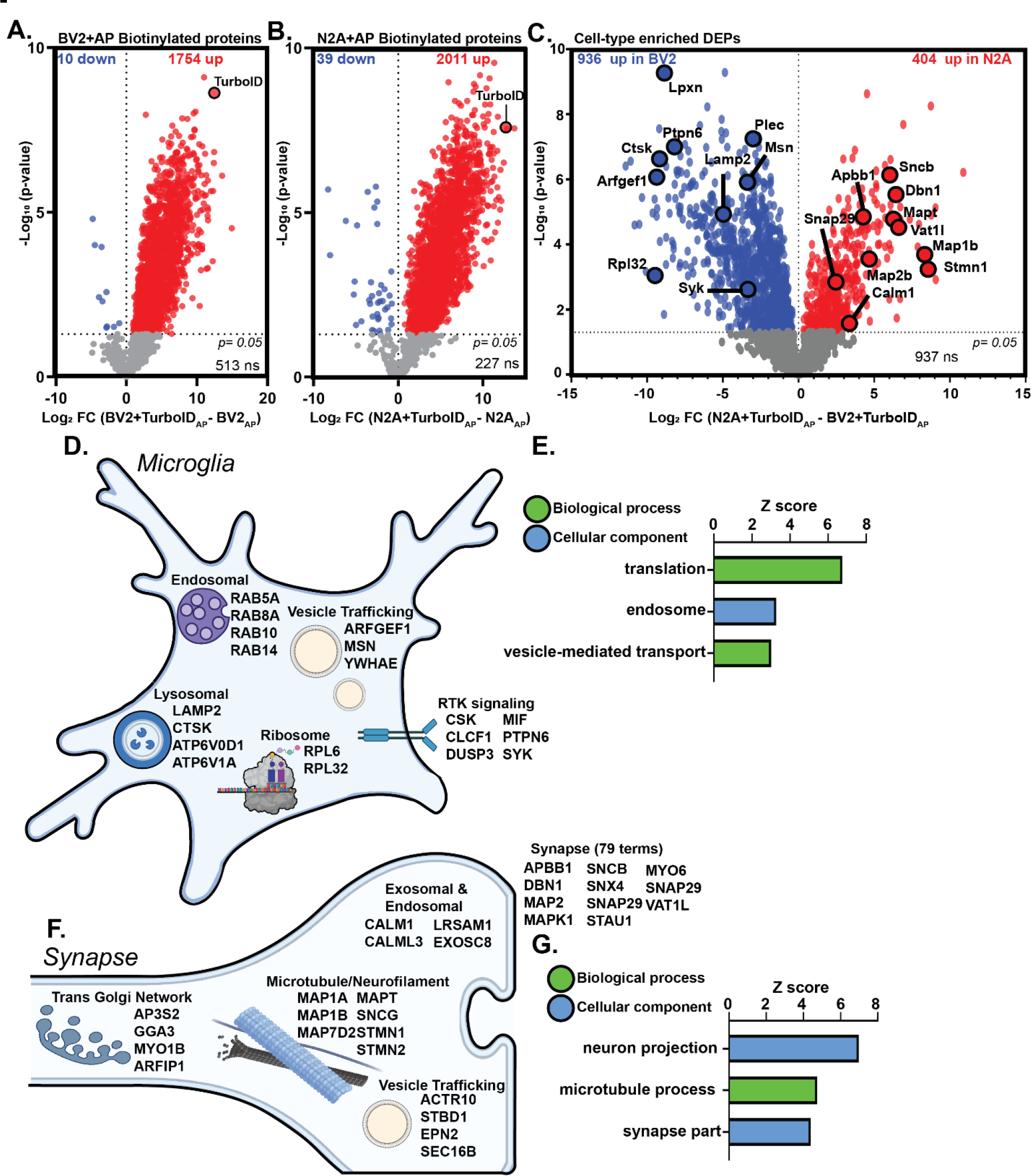
TurboID labeling and Streptavidin AP captures cellularly distinct proteomes. **A.** DEA of BV2 AP samples showing robust TurboID biotinylation of over >1700 proteins over endogenously biotinylated proteins derived from untransduced cell lines. **B.** DEA depicting biotin enrichment of N2A transduced AP proteome reveals >2000 proteins labeled by TurboID **C.** DEA comparing transduced AP proteomes of BV2 (left) and N2A (right) TurboID-biotinylated proteomes. There are 936 proteins labeled by TurboID enriched in BV2 and 404 proteins enriched in N2A biotin-labeled proteomes. Proteins with disease-relevance to neurodegenerative disease are highlighted. **D.** Schematic of the variety of subcellular compartments labeled by TurboID in microglia (BV2). **E.** Gene Ontology (GO) of highly enriched cellular components and biological processes within the biotin-labeled BV2 proteome. TurboID biotinylates translational machinery, endosomal machinery, and vesicle-bound membranes in BV2 cells. **F.** Schematic of diversity of subcellular compartments labeled by TurboID in synaptic compartment (N2A). **G.** GO of significantly enriched cellular components and biological processes within the biotin-labeled N2A proteome confirms that TurboID biotinylates neuronal processes including synaptic machinery and neuron projection.

To inform the application of CIBOP to future mouse models of disease, we tested the hypothesis that global cytosolic biotinylation of TurboID-NES can achieve a proteomic breadth sufficient for conclusive cellular distinction between glia and neurons. We performed DEA on AP proteomes from BV2 and N2A cell lines stably expressing TurboID- NES (N2A+TurboID-NES_AP_ – BV2+TurboID-NES_AP_). We identified 936 proteins enriched in BV2+TurboID-NES AP samples and 404 proteins enriched in N2A+TurboID-NES AP (**Fig 5C; SD 2G**). Notably, TurboID-NES biotinylated proteins with relevance to Alzheimer’s disease (AD) in both microglial and neuronal cells. For example, Moesin (Msn), protein tyrosine phosphatase nonreceptor 6 (Ptpn6), C-terminal Src Kinase (Csk) and Plectin (Plec) were highly enriched in microglial proteomes labeled by TurboID-NES. Moesin has been identified as a hub protein for AD etiology in both human and 5xFAD mouse models ^29–32^ and is necessary for P2X7R-dependent proteolytic processing of amyloid precursor protein^33^. Both Csk and Ptpn6 have been identified as AD hub genes.^34^ Ptpn6 is associated via signaling pathways with CD33, a risk locus identified in human AD genome wide association studies.^34–37^ In N2A cells, TurboID-NES biotinylated AD-relevant proteins including Microtubule-Associated Protein tau (Mapt), Microtubule-associated protein 1A/B (Map1a; Map1b), amyloid beta precursor protein binding family B (Apbb1), and Calmodulin 1 (Calm1).

Without being directed to a specific subcellular compartment, TurboID-NES biotinylated a variety of proteins in microglial and neuronal cell lines. In BV2 cells, TurboID- NES biotinylated lysosomal, ribosomal, endosomal, and vesicular proteins. Additionally, TurboID-NES biotinylates proteins important to receptor tyrosine kinase signaling (RTK) (**Fig 5D**). Gene-Set-enrichment analysis (GSEA) of microglial-enriched proteins biotinylated by TurboID-NES highlighted significant enrichment for translational machinery, endosomal proteins as well as intracellular trafficking of vesicles (**Fig 5E**). In N2A cells, TurboID-NES labeled proteins involved with trans-Golgi network trafficking, microtubule and neurofilament elements, endosomal and exosomal machinery, vesicular trafficking, and over 75 synaptic terms out of 404 terms identified in the enriched N2A biotinylated proteome (**Fig 5F**). GSEA of the N2A biotinylated proteome identified top terms associated with neuron projection, microtubule processes, and synaptic parts, supporting the ability of TurboID-NES to label synaptic proteins which could contribute to cell-type distinction (**Fig 5G**). Using the proteome of cellularly distinct proteins labeled by TurboID-NES identified in the DEA in Figure 4A, we mapped proteins to risk-loci associated with neurodegenerative diseases (**SF 3**). Using risk loci published in the Alzheimer’s disease (AD) MAGMA^6^, Parkinson’s disease (PD) MAGMA^38^, and Amyotrophic lateral sclerosis / Frontotemporal dementia (ALS/FTD) MAGMA^39^ datasets, we identified 82 and 46 AD-relevant and cellularly distinct proteins labeled by TurboID-NES in BV2 and N2A proteomes, respectively (**SF 3A**). We identified 32 and 17 PD-relevant and 84 and 29 ALS/FTD-relevant proteins in BV2 and N2A proteomes labeled by TurboID-NES and enriched by streptavidin-based affinity purification (**SF 3B, 3C**). Taken together, TurboID-NES, when directed into the cytosol for global biotinylation of proteins, biotinylated a breadth of proteins sufficient to distinguish between microglial and neuroblastoma cell lines. Additionally, TurboID-NES biotinylated proteins critical to neurodegenerative etiologies. These proof-of-principle analyses support the future direction of TurboID-NES into distinct brain cell types within living mouse models of neurodegeneration.

### BV2 proteomes biotinylated by TurboID-NES capture Lipopolysaccharide driven changes, partially reflected in the whole-cell BV2 proteomes

After confirming that TurboID-NES robustly labeled distinct cellular proteomes with minimal impact on homeostatic phenotype, we hypothesized that TurboID-NES could label proteins impacted by LPS treatment. Our dimension reduction analyses confirmed our ability to resolve proteomic differences in transduced AP cells based on cell type and LPS challenge (**SF 1B**). To understand global differences between WC and AP proteomes differentially expressed by LPS treatment, we created a heatmap representation of LFQ intensity values (2350 proteins; **SD 2H**). We identified 6 distinct proteomic clusters (**Fig 6A**), as determined by the elbow-method optimization of number of clusters (**SF 4**). Two large clusters (Cluster 1 and Cluster 2) showed specific abundance differenced based on affinity purification (AP) status. Cluster 1 (943 proteins) was highly abundant in whole cell lysates as compared to AP samples with minimal effect of LPS, and was enriched in nuclear, mitochondrial, RNA binding and metabolic proteins (**Fig 6B, 6C**); proteins not readily accessible by a cytosolic-directed TurboID-NES. Cluster 2 (608 proteins) was conversely more abundant in BV2 AP proteins compared to whole cell lysates with minimal effect of LPS, and were enriched in cellular membrane organization, endocytic, cytoplasmic location, RNA transport, ER lumen, translation and vesicle-mediated transport functions, consistent with groups of proteins preferentially biotinylated by TurboID-NES. More importantly, we identified clusters of proteins that captured LPS effects in WC lysates, or AP samples. LPS significantly decreased protein abundances in BV2 AP in clusters 3 (191 proteins) and 4 (236 proteins) and significantly increased protein abundances 5 (131 proteins) and 6 (241 proteins) (**SF 5**). Cluster 3 was enriched in cytosolic, lysosomal (hydrolase activity), and ribosome proteins, suggesting that the TurboID-NES approach captured an effect of LPS on translation and lysosomal functions that cannot be resolved at the whole cell level. Cluster 5, the ‘BV2+ LPS shared cluster’, was increased by LPS treatment in both WC and AP samples, and showed enrichment in terms such as response to IFN-γ, oxidative stress (hydrogen peroxide related processes), and peroxisome and phagosome functions, indicative of expected LPS-driven proteomic changes in microglia. Several inflammatory terms in Cluster 5 were previously reported in BV2 cells specifically in response to 1 µg/mL LPS for 48 hours ^40^, which included antigen processing and presentation, and glycolysis / gluconeogenesis, reflecting a shift in bioenergetics induced by inflammatory stimuli (**Fig 6B, 6C**). Cluster 6 showed LPS- induced increased levels that were only apparent in AP samples, and were enriched in RNA binding, nucleolus localization, ribosome, translational activity and spliceosome functions. Cluster 6 may represent altered localization of RNA-interacting splicing proteins, as well as potential nuclear speckle and nucleolar proteins from the nucleus to the cytoplasm due to LPS-induced stress. Such altered localization events are more likely to be captured using the TurboID-NES approach, rather than at the whole cell level. To rule out the possibility that LPS impacts TurboID-NES localization in BV2 cells, we performed ICC and WB studies on cytoplasmic and nuclear fractions (**SF 6A,6B,6E**). We performed colocalization analyses comparing the DAPI signal as a nuclear marker with V5 for TurboID-NES or StrepDylight to determine the percent of cytosolic TurboID-NES signal or biotinylation signal, respectively. Interestingly, LPS significantly increased the percent of cytosolic TurboID-NES and cytosolic biotinylation (**SF 6C,6D**). We observed predominantly cytosolic localization of TurboID-NES (via V5 localization) and biotinylation in both ICC studies, as well as in WB analyses (**SF 6**). Taken together, it is likely that cytosolic direction of TurboID-NES can identify aberrantly trafficked proteins in response to LPS, though further studies are necessary.

**Figure 6.**
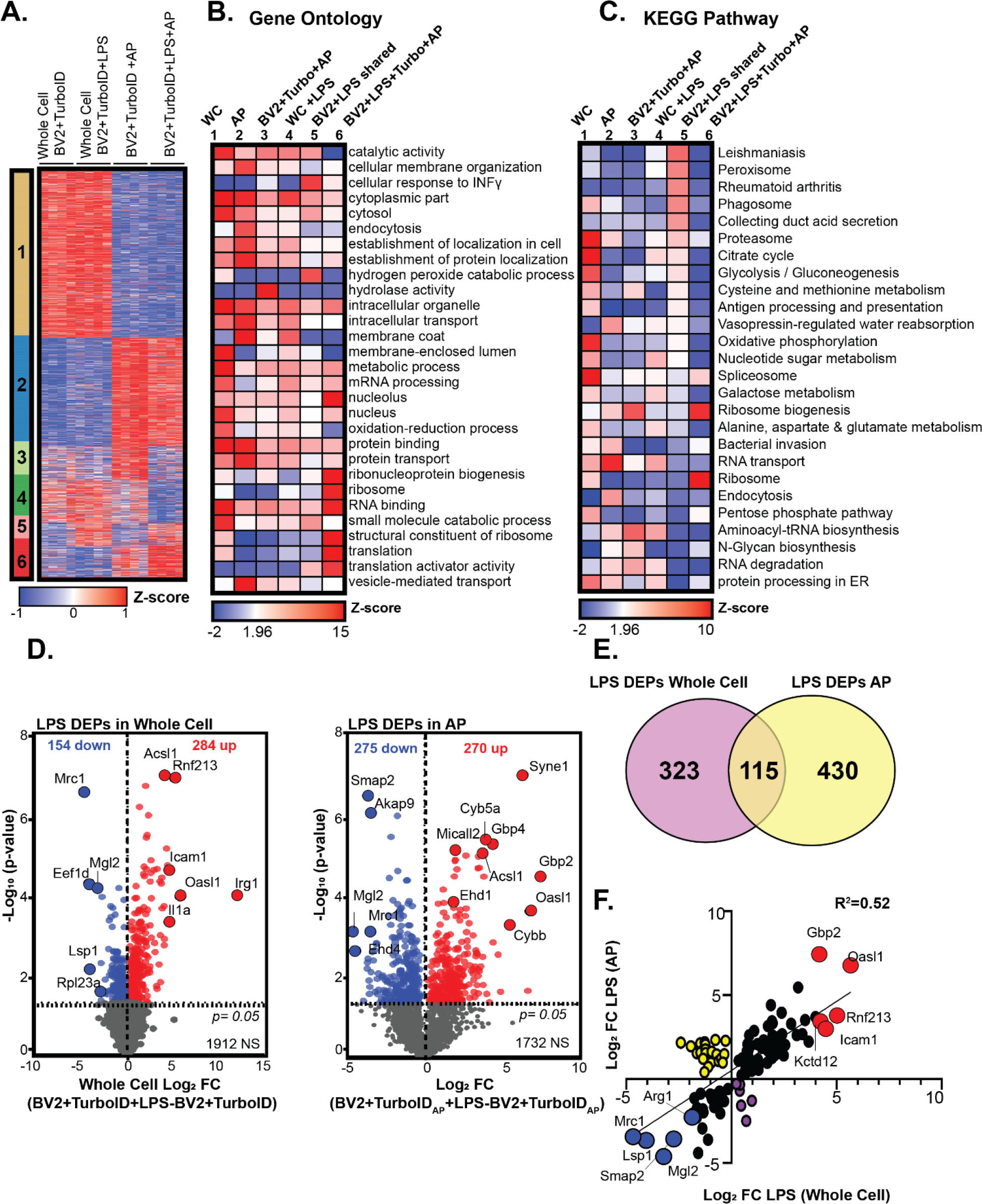
TurboID-mediated biotinylation partially captures LPS-driven proteomic changes. **A.** K-means clustered heatmap representation of LFQ intensity data of proteins differentially expressed by TurboID in whole-cell and transduced AP BV2 proteomes (n = 4 per experimental group). **B.** Heatmap representation and functional annotation of gene ontology from *murine* GO database. Heatmap color intensity depicts Z-score values. **C.** Heatmap representation of z-scores associated with LPS DEPs derived from KEGG pathways database. **D.** DEA of LPS DEPs in WC (*left*) and AP (*right*) identifies 438 and 545 LPS DEPs, respectively. **E.** Venn diagram depicting 323 LPS DEPs unique to whole-cell BV2 samples, 115 shared LPS DEPs and 430 unique LPS DEPs in BV2 AP samples. **F.** Linear regression of Log2FC values of BV2 AP samples (y-axis) and BV2 whole-cell samples (x-axis) reveals a modest correlation (r^2^=0.52) of log2FC of the shared 115 LPS DEPs. Red points depict proteins the top 5 proteins significantly upregulated with LPS treatment in AP and whole-cell samples. Yellow points represent proteins significantly increased in AP samples and significantly decreased in whole-cell samples with LPS treatment. Blue points illustrate the top 5 proteins significantly decreased with LPS in both AP and whole-cell BV2 samples. Purple points depict proteins which are significantly increased in the whole-cell proteome and significantly decreased in the AP proteome with LPS treatment.

We then compared the impact of LPS treatment on WC and AP proteomes using DEA. DEA of proteomic differences induced by LPS treatment in BV2 WC samples (BV2+LPS+TurboID-NES_WC_ – BV2+TurboID-NES_WC_) and AP samples (BV2+LPS+TurboID-NES_AP_ – BV2+TurboID-NES_AP_) identified 438 proteins impacted by LPS in the WC proteome and 535 proteins impacted by LPS in AP samples (**Fig 6D; SD 2J, SD 2I**). The top proteins increased with LPS treatment in the WC BV2 proteome included Immune-responsive gene 1 (IRG1), oligoadenylate synthetase-like 1 (Oasl1), interleukin 1 a (Il1a), Ring Finger Protein 213 (Rnf213), long-chain acyl-CoA synthetase family member 1 (Ascl1), and intracellular adhesion molecule 1(Icam1). The proteins most down-regulated by LPS treatment in BV2 WC samples included Macrophage Mannose Receptor 1-Like Protein (Mrc1) macrophage galactose N-acetyl-galactosamine specific lectin 2 (Mgl2), and Eukaryotic translation elongation factor 1 delta (Eef1d) (**Fig 6D**). The top terms increased with LPS in BV2 AP samples included proteins such as Oasl1, Gbp2, Syne2, Cyb5a, and Acsl1. Proteins in the BV2 AP proteome most down-regulated by LPS treatment included Mgl2, Smap2, and Mrc2. LPS-increased proteins shared between the WC and AP samples, including Il1a, Irg1, Oasl1, corresponded with pro-inflammatory M1 phenotypic markers, as well as non-canonical inflammasome mediators such as Gbp2.^41–46^

Proteins similarly decreased with LPS treatment shared between WC and AP BV2 proteins including canonical M2 markers such as Arg1 and Mgl2.^47–50^ When comparing the overlap of proteins differentially expressed by LPS treatment in the WC and transduced AP BV2 proteomes, we identified 115 shared proteins that were differentially expressed in both AP and WC BV2. 323 proteins were differentially expressed in response to LPS only in the WC proteome while 430 proteins were differentially expressed in response to LPS only in the AP proteome (**Fig 6E**). Of the 115 proteins with shared LPS-induced differential expression we observed a moderate concordance based on magnitude (Log2FC LPS _WC_ Vs. Log2FC LPS _AP_) and direction of LPS effect (Coefficient of determination, R^2^= 0.52) (**Fig 6F**). We identified 28 proteins that showed incongruent changes (**yellow points, Fig 6F**) with LPS-induced increase in AP proteomes but LPS-induced decrease in WC proteomes.

## 4. DISCUSSION

Proteomics based on proximity-labeling strategies using biotin ligases (e.g. BioID, TurboID-NES) are being increasingly used in *in vitro* and *in vivo* experimental contexts and in model systems ranging from plants^17,51–53^, yeast^54^, zebrafish^55,56^, Drosophila^57,58^, *C. elegans*^16,59^, to mouse models^18,21,22,60^. TurboID is one of the most efficient biotin ligases that can effectively label several proteins within a 10nm labeling radius in mammalian cells. The high catalytic activity, non-toxicity, and promiscuity of TurboID uniquely position TurboID as a powerful tool to obtain global proteomic snapshots of specific cells in homeostatic and disease states, particularly in multi-cellular models. Many recent studies have incorporated TurboID and split-TurboID^20^ to characterize interactomes and secretomes via fusion with proteins, organelles, and inter-cellular contacts. ^14,16,21,57, 60–64^ While one application of TurboID-based proteomics is to identify protein-protein interactors of proteins of interest or within specific subcellular compartments, another application of TurboID is to broadly label the cellular proteome of a specific cell type, so that cell type-specific proteomics can be resolved from a complex mixture of proteins derived from multiple cell types. The viability of the latter application was recently tested *in vivo* using genetic Cre/lox strategies to resolve neuronal and astrocyte proteomes in the native state of these cells in mouse brain, a method referred to as cell type-specific *in vivo* biotinylation of proteins (CIBOP).^22^ Whether these neuronal or glial proteomes obtained using CIBOP reflect global cellular proteomes, remains to be clarified. As interest in TurboID-based global cellular proteomics continues to grow, it is important to determine what fraction of the whole cell proteome in mammalian cells can be faithfully captured by the TurboID-NES approach, under both homeostatic (resting) conditions and following cellular perturbations (eg. immune stress by LPS to mimic neuroinflammatory disease conditions). It is also important to determine whether TurboID-NES over-expression or excessive biotinylation impacts molecular phenotypes and cellular functions, and whether TurboID-NES- biotinylated proteomes have inherent biases as compared to the whole cell proteome.

These questions can be answered by performing well-controlled *in vitro* studies using distinct types of mammalian cells. The results from such studies can inform the interpretation of proteomic findings gained from *in vitro* and *in vivo* applications of TurboID- NES in mammalian model systems, while considering the relative biases of this strategy for cell type-specific proteomics. To address these questions, we directed TurboID-NES to the cytosol using a NES, rather than using protein-specific or cellular compartment-restricted localization of TurboID-NES. We hypothesized that TurboID-NES could globally biotinylate a breadth of the cellular proteome sufficient to distinguish two distinct brain cell types (neurons and microglia) and capture the effect of an inflammatory challenge, without significantly impacting functional or molecular phenotypes of cells that over-express TurboID-NES.

We generated murine N2A neuroblastoma and BV2 microglial cell lines that stably express TurboID-NES, and validated the expression and functionality of TurboID-NES using flow cytometry, biochemical, and immunocytochemical analyses. After validation of these stably-transduced TurboID-NES cell lines, we analyzed the TurboID-NES- biotinylated as well as the WC reference proteomes of sham-treated or LPS-treated cells using MS-based quantitative proteomics. We confirmed that TurboID-NES biotinylates >50% of the whole cell proteome in both N2A and BV2 cells, including proteins in a variety of subcellular compartments in the cytosol (e.g., endocytic machinery, ribosomal proteins, mRNA binding proteins, membrane proteins, vesicle-related and transport proteins and cytoskeletal proteins). While a large proportion of biotinylated proteins were common to both cell types, several neuron-enriched and microglia-enriched proteins were indeed identified in the respective biotinylated proteomes. For example, BV2 microglial biotinylated proteomes captured endo-lysosomal and phagocytic proteins while neuron-projection and axonal transport proteins were labeled in N2A neurons, further verifying the validity of this approach to study cell type-specific mechanisms of distinct cell types rather than just homeostatic cellular mechanisms that are shared across cell types. These neuron-enriched and microglia-enriched proteins captured by the TurboID-NES approach also included several neurodegenerative disease-related proteins with causal implications in AD, PD and ALS. This suggests that the TurboID-NES approach can be used to investigate disease-relevant biology in neurons and glia in mammalian systems.

Using the TurboID-NES approach, we also captured immune effects of LPS on BV2 cells. Some of these were shared at the whole cell level (eg. increased expression of pro-inflammatory proteins Gbp2, Oasl1, Rnf213 and Icam1 and decreased expression of anti-inflammatory proteins such as Mrc1, Arg1 and Mgl2), indicating that TurboID-NES biotinylation captures core pathological transformations induced by long-term LPS stimulation in BV2 cells. KEGG terms in this cluster include antigen processing and presentation, glycolysis and gluconeogenesis, and alanine, aspartate and glutamate metabolism. ^40^ Previous studies assessing the longitudinal impact of LPS stimulation on BV2 proteomes similarly identified these major KEGG terms reflecting a strong metabolic shift in later LPS activation. Our results confirm that TurboID-NES expression does not impair metabolic functioning, and we confirm that biotinylation by TurboID-NES is able to biotinylate metabolic and immune responsive protein pathways impacted by long-term LPS stimulation. Importantly, LPS DEPs identified in transduced and affinity-purified BV2 cells correlate moderately with LPS DEPs at the WC level. Despite these consistent LPS- induced proteomic changes observed in WC and biotinylated BV2 proteomes, a relatively large group of protein changes due to LPS were identified only at the level of biotinylated proteins but not at the whole cell level. These LPS-induced proteomic changes preferentially captured by the TurboID-NES approach may be due to better access of TurboID-NES to specific cellular compartments, such as the cytosol, which allow capture of post-translational effects of LPS, such as altered protein trafficking, nucleocytoplasmic transport of proteins from the nucleus to the cytosol or vice versa and altered localization of RNA binding, ribosomal and ribonucleoprotein-related proteins. Consistent with this, proteins involved in RNA binding, nucleolar proteins, ribosomal and translational machinery were selectively increased in the LPS-induced biotinylated proteome. We also observed a slight increase in trafficking of TurboID-NES to the cytosol with LPS stimulation, implying that the effects of LPS observed at the level of the biotinylated proteome are likely due to a combination of increased cytosolic localization of TurboID- NES and altered localization of biotinylated proteins. The LPS-induced changes in levels of ribonuclear proteins agree with reported ribosomal mechanisms involved in innate immune activation in microglia, that are responsible for translational repression and a divergence between mRNA and protein expression following LPS challenge.^65^ Furthermore, ribonuclear proteins within nuclear speckles may also be localized to the cytosol as a direct result of immune activation or a cell proliferative response to LPS stimulation, as has been reported previously.^66,67^ Finally, several nucleus-resident RNA binding and ribonucleoproteins can translocate to the cytosol to regulate mRNA translation by the ribosome. The ability to use TurboID-NES-based proteomics to investigate protein trafficking and mis-localization is of particular relevance to neurodegenerative disorders. Mis-localization of nuclear proteins to the cytosol occur in several neurodegenerative diseases, where RNA-binding proteins such as TDP-43, Tau and FUS can aberrantly localize to the cytosol where they become more prone to aggregation.^68^ Therefore, the TurboID-NES approach when employed *in vivo* via CIBOP, could be specifically used to interrogate mechanisms of neurodegeneration that involve dysfunction of nucleocytoplasmic transport, changes in protein trafficking, and cytosolic aggregation.

Another important result of our study is that TurboID-NES over-expression had a minimal impact on the proteomes of both N2A and BV2 cells at the whole cell level and did not significantly impact cellular respiration or secreted cytokine profiles in response to LPS in BV2 cells. These findings suggest that cellular molecular composition and function are not meaningfully compromised using the TurboID-NES approach. This finding is consistent with the lack of electrophysiological alterations observed in *in vivo* TurboID-NES studies in Camk2a excitatory neurons.^22^ These findings are indeed reassuring and support the use of TurboID-NES-based cellular proteomic profiling approaches to investigate mechanisms of disease with minimal effects on cellular functions due to TurboID-NES over-expression.

Despite the strengths of well-controlled *in vitro* studies, some limitations of our work need to be considered. Cell lines such as BV2 and N2A cells, despite their ability to recapitulate major cellular phenotypes of microglia and neuronal cells, display many well-known differences as compared to primary cells in the nervous system. Another limitation is that the LPS dose and duration used for (1 µg/mL for 48 hours) could have caused induced over-stimulation, cell-death, apoptosis or stress-induced changes in BV2 cells, leading to biased proteomic findings. However, it is reassuring that our observed LPS effects on the BV2 whole cell proteome, generally agree with prior proteomics studies of BV2 cells using low dose (10-100ng/mL) and high concentrations (1 µg/mL).^69,70^ Another limitation is related to the use of the NES in our TurboID studies. TurboID-NES was intentionally directed to the cytosol to increase the sampling of non-nuclear proteins, as well as to minimize undesired effects of excessive biotinylation of nuclear or mitochondrial proteins that are involved in key cellular functions, such as chromosomal stability and gene regulation and mitochondrial metabolic processes. However, in doing so, we also biased the biotinylated proteome away from nuclear, mitochondrial and intraluminal-directed proteins, which was evident when biotinylated AP proteomes were contrasted with WC lysate proteomes. Future studies may consider comparing proteomes using TurboID without NES to TurboID-NES to confirm whether cellular toxicity is indeed observed. To minimize the chance of relative biotin deficiency due to TurboID-NES over-expression, all experimental conditions included biotin supplementation in the medium. While this controlled for biotin deficiency, this may have induced some metabolic alterations due to excessive biotin in experimental conditions.

Our results demonstrate the ability of TurboID-NES-based cellular proteomics to capture a representative portion of disease-relevant and immune-relevant proteins in two distinct brain cell types, namely microglia and neurons, using immortalized cell lines. These results directly impact future directions of TurboID-NES using the CIBOP approach in transgenic mouse models of inflammation and neurodegeneration.

## 5. CONCLUSIONS

In conclusion, we generated transduced neuroblastoma and microglial cell lines expressing cytosolic TurboID-NES which yielded robustly labeled proteomes that covered a wide variety of subcellular compartments with no significant impact to cellular phenotypes. We identified a high representation of neurodegenerative disease-relevant protein pathways as well as a partial coverage of immune-relevant proteins in microglia. The breadth of the proteome labeled by TurboID-NES distinguished neuroblastoma cells from microglial cells, and TurboID-NES labeled over 50% of identified proteins within each cell type; supporting the *in-vivo* application of TurboID-NES in its ability to purify cellularly-distinct proteomes. Our results also highlight inherent biases of TurboID-NES-based proteomics approaches which may be more suited to investigate post-translational mechanisms such as protein trafficking which are not captured by whole cell proteomics.

## 6. METHODS

### 6.1 Antibodies, Buffers & Reagents

A complete table of antibodies & reagents are provided (**Tables 1 & 2**).

**Table 1.**
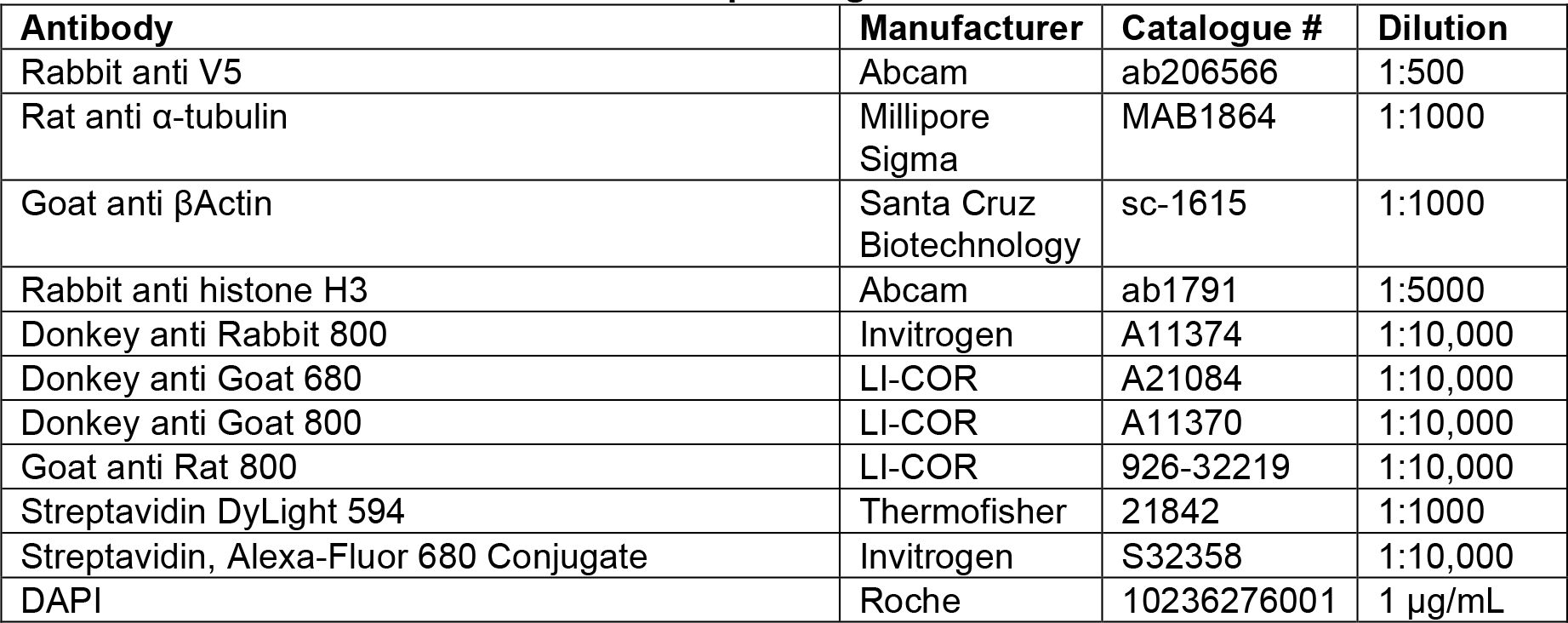
Antibodies used and their corresponding dilutions.

| Antibody | Manufacturer | Catalogue # | Dilution |
| --- | --- | --- | --- |
| Rabbit anti V5 | Abcam | ab206566 | 1:500 |
| Rat anti α-tubulin | Millipore<br>Sigma | MAB1864 | 1:1000 |
| Goat anti βActin | Santa Cruz<br>Biotechnology | sc-1615 | 1:1000 |
| Rabbit anti histone H3 | Abcam | ab1791 | 1:5000 |
| Donkey anti Rabbit 800 | Invitrogen | A11374 | 1:10,000 |
| Donkey anti Goat 680 | LI-COR | A21084 | 1:10,000 |
| Donkey anti Goat 800 | LI-COR | A11370 | 1:10,000 |
| Goat anti Rat 800 | LI-COR | 926-32219 | 1:10,000 |
| Streptavidin DyLight 594 | Thermofisher | 21842 | 1:1000 |
| Streptavidin, Alexa-Fluor 680 Conjugate | Invitrogen | S32358 | 1:10,000 |
| DAPI | Roche | 10236276001 | 1 μg/mL |

**Table 2.**
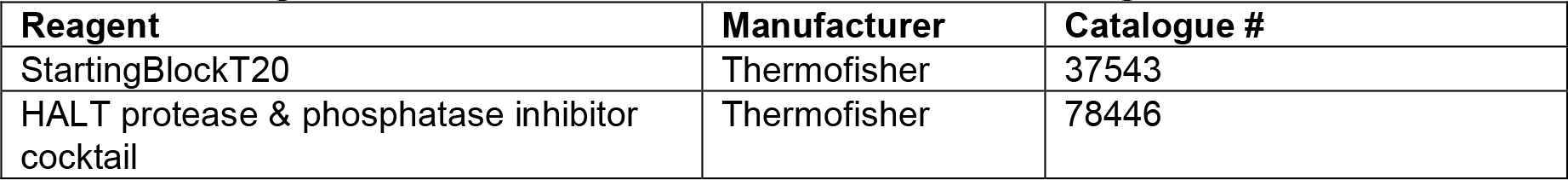
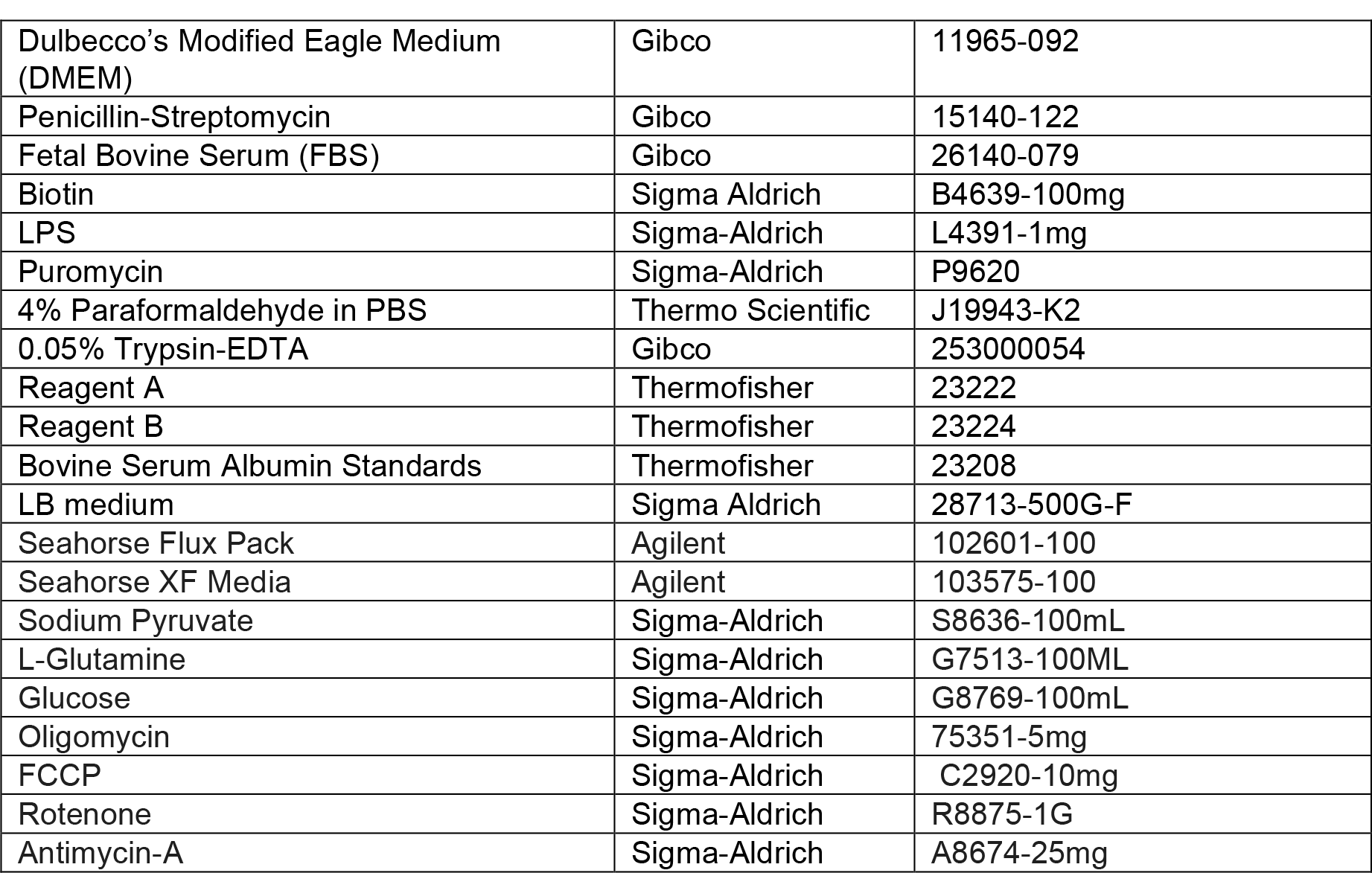
Reagents used and their manufacturer and catalogue numbers.

### 6.2 Cell Culture

N2A and BV2 cells were cultured in filtered Dulbecco’s Modified Eagle Medium (DMEM) supplemented with high glucose and L-glutamine containing 1% penicillin-streptomycin, and 10% Fetal Bovine Serum (FBS). All media was vacuum-filtered with 0.2 µm filters. The cells incubated at 37 degrees Celsius (°C) and 5% CO2 until reaching 80% confluency. The splitting regimen took place twice weekly, plating one million cells onto a 100mm culture plate to a final volume of 10mL culture media. In preparation for Mass Spectrometry (MS) experiments, cells reached 95% confluency in 150 mm plates.

Transduced cells expressing TurboID-NES were kept in 2 µg/mL puromycin media until being plated for MS, wherein they forwent puromycin treatment.

### 6.3 Genetic Constructs & Gene Delivery

The V5-TurboID-NES_pCDNA3 and plasmid is a gift from Alice Ting (Addgene plasmid #107169). Plasmids were transformed using a competent *E. coli* strain DH5α according to manufacture protocols. Briefly, DH5 α cells were thawed on ice before aliquoting 50 µL into 1.5mL tubes. Constructs were diluted 1:1000 into autoclaved milli-Q water. LB medium was prepared and autoclaved by diluting 20 grams of LB Broth, Vegitone into 1000mL of H2O. To the 50ul aliquots of DH5α competent cells, 5 µL of diluted constructs were mixed by turning the tubes upside down. The plasmids were incubated with the DH5α competent cells for 30 minutes on ice. Following incubation, the samples underwent heat-shock by for 42°C for 20 seconds and were placed on ice for 2 minutes. 500 µL of prewarmed LB medium were added to each sample before being placed on a rotor to shake at 225 rpm at 37°C for 1 hour. Plasmid DNAs were purified using QIAfilter Plasmid kits (Midi pre kit, Qiagen #12243) following manufacturer protocol. Restriction sites (underlined) were introduced via the following polymerase chain reaction (PCR) primers (V5.bstb.S; 5’- gcgcctactctagagctagcgaa<u>ttcgaa</u>gccaccatgggcaagcccatccccaa-3’) (nes.Bam.A; 5’- agaaggcacagtcggcggccgcg<u>gatcc</u>ttagtccagggtcaggcgctccagggg-3’). The V5-TurboID-NES sequence was subcloned into pCDH-EF1-MCS-BGH-PGK-GFP-T2A-Puro (CD550A-1) and sequenced. N2A, and BV2 cells were transduced with a puromycin-resistant lentivirus construct of a V5-TurboID-NES containing a nuclear export sequence (V5-TurboID-NES) and a green fluorescence protein (GFP) connected via a T2A linker. Constructs were generated in Emory University’s Viral Vector Core. Given a titer of 1.5x10^9^ I.U./ml, we experimented with multiplicity of infection (MOIs) 5 and 10. In biological triplicates, 3 wells of each cell type received the lentivirus at either a MOI of 5 or 10, or received no virus (untransduced control). 48 hours following transduction, ½ of the media was replaced with fresh media, and the cells were split 1:3 on 72 hours following transduction. Puromycin selection began 96 hours after transduction, ½ of the media was replaced with 2 µg/ml of puromycin for a final concentration of 1 µg/ml. For the following week, ½ of the media was replaced every other day with fresh puromycin-containing media, to remove non-adhering cells. After this, the cells were split twice weekly and maintained with media containing 1 µg/ml of puromycin. We validated the puromycin screening procedure by assessing the percentage of GFP positive cells with a fluorescent microscope. A majority of cells were GFP positive three days after addition of puromycin selection. After >90% of the cells were GFP positive, the cells were maintained in 10 cm dishes in media containing 1 µg/ml of puromycin. Cells receiving the lentivirus at a MOI of 5 reached confluency sooner, and we did not observe a difference in transduction efficacy between MOI’s 5 and 10, which were subsequently pooled.

### 6.4 Generation of whole cell lysates and supernatants and Confirmation of labeling

BV2 and N2A cells transduced or untransduced with V5-TurboID-NES were seeded onto 10cm plates. 24 hours after plating, media was replaced either with biotin-supplemented media (200μM) or media containing both LPS (1μg/mL) and biotin supplementation (n = 6/group). 48 hours after media replacement, the media was taken off, centrifuged for 2 minutes at 800 rpm room temperature to remove cellular debris, and preserved in a 15mL tube. Cells were dissociated using 2.5mL of 0.05% trypsin-EDTA. After trypsin dissociation, cells were rinsed and collected via manually pipetting with 7.5mL fresh media before being transferred to 15mL tubes. Supernatants and cell pellets were flash-frozen on dry-ice. Dissociated cells and supernatants were centrifuged at 800rpm for 2 minutes (RT). Supernatants were transferred to a fresh 15mL tube before being flash-frozen on dry ice. Cell pellets were washed twice with 10mL of ice-cold 1x phosphate buffer saline (PBS). Finally, cell pellets were resuspended in 1mL of ice-cold PBS and transferred to 2 mL LoBind Eppendorf tubes (Eppendorf #022431102). Cells were centrifuged for 2 minutes at 800rpm before being flash-frozen on dry ice. Cell pellets were harvested in 500μL Urea lysis buffer (8M Urea, 10mM Tris, 100mM NaH_2_PO_4_, pH 8.5) with 1x HALT protease & phosphatase inhibitor cocktail without EDTA. Cell lysates were then sonicated at 30% amplitude thrice in 5-second on-off pulses to disrupt nucleic acids and cell membrane. All cell lysates were centrifuged at 4°C for 2 minutes at 12,700rpm. The supernatants were transferred to a fresh 1.5mL LoBind Eppendorf tube. The protein concentrations of whole cell lysates were determined by Bicinchoninic acid (BCA) assay reagents using Bovine Serum Albumin Standards. Out of the 6 samples per experimental group prepared, 4 samples per group were reserved for quality-control analyses performed prior and in addition to MS studies.

### 6.5 Immunoblotting

In each well, 10-20 µg of protein from cell lysates resolved in a 4-12% polyacrylamide gel and transferred onto iBlot 2 Transfer Stack containing nitrocellulose membrane using the BOLT transfer system. The membranes incubated for 1 hour at room temperature in StartingBlockT20 before receiving rabbit anti V5 primary antibody overnight at 4 °C or 1 hour at room temperature. After primary antibody incubation, the membranes underwent three rapid washes with 1x TBST followed by three 10-minute washes with 1xTBST. Membranes then underwent three rapid washes with 1x TBS and three 10-minute washes with 1xTBS. The membranes incubated for 1 hour at room temperature in a secondary antibody cocktail of streptavidin 680 to visualize biotinylated proteins and donkey anti rabbit 800 to visualize V5- tagged TurboID-NES. The membranes were then washed again as previously described before undergoing imaging via the Odyssey infrared Imaging System (LI-COR Biosciences).

### 6.6 Immunofluorescence

Immunofluorescent staining was performed as published previously with modifications for cultured cells.^22^ Briefly, BV2 and N2A cells were seeded at ∼50,000 cells onto autoclaved and HCl-ethanol-treated 25mm coverslips in a 6-well dish. All cells received supplemental biotin treatment (200μM) for the duration of maintenance. After reaching 50% confluency, cells were washed thrice with warm sterile PBS to remove media. Cells were fixed with 4% paraformaldehyde in PBS for 30 minutes and washed thrice in ice-cold PBS for 10 minutes, gently orbitally-rotating at 50 rpm (IKA, # KS 260). Fixed-cells were permeabilized and blocked simultaneously in a solution of 5% normal horse serum (NHS) in 0.25%TBST diluted in PBS for 1 hour at room temperature on the orbital-rotator. To each well, cells received 1mL of rabbit anti V5 (1:500) primary antibody solution in 1% NHS diluted in PBS overnight at 4 °C. All cells received a 5-minute incubation with DAPI for nuclear staining. All IF imaging was performed with a 60x oil-immersion objective taken using Keyence BZ-X810. Colocalization analysis was performed with slight modifications from previous methods.^71,72^ We subtracted the target area (µm^2^) of DAPI-signal overlap with V5 or biotinylation signal (nuclear-localizing TurboID-NES or biotinylated proteins, respectively) from the total area of V5 or Streptavidin signal using the Keyence BZ-X810 Analyzer hybrid-cell-count colocalization software. Only cells positive for both DAPI and V5 or Streptavidin DyLight were analyzed for colocalization analysis. Significance p values were assessed using the two-tailed Mann-Whitney test. The total target area counts (number of cells counted due to the presence of a cell with overlapping of DAPI and V5 or Streptavidin DyLight signal) for transduced BV2 cells assessed for V5/DAPI colocalization was 1876 cells. The total target area counts for V5/DAPI colocalization for transduced BV2 cells receiving LPS challenge was 1945 cells. The total Streptavidin DyLight/DAPI colocalization target counts for transduced naïve BV2 cells was 2289 cells. The total target counts for Streptavidin DyLight/DAPI colocalization for transduced BV2 cells receiving LPS was 1638 cells.

### 6.7 Subcellular fractionation

Transduced and untransduced BV2 and N2A cells were plated in triplicates on 10cm plates and grown to 70% confluency. Cellular monolayers were rinsed with 10mL ice cold PBS, and PBS was aspirated. Cell monolayers were scraped with a sterile cell scraper in 2.97 mL PBS with 30 µL of 100x HALT, added immediately prior to use. Cell slurries were transferred to 15mL tubes and centrifuged at 1,000 x g at 4°C for 5 minutes. The supernatants were removed and the pellet was washed once in 1 mL cold PBS. To obtain the WC fraction, 100 µL of the cell slurry was transferred to a fresh 0.5 mL Eppendorf tube. The remaining 900 µL of cell slurry was centrifuged at 1,000 x g at 4°C for 5 minutes. Supernatants were removed and 150 µL of Hypotonic Lysis buffer (10 mM HEPES, pH 7.9, 20 mM KCl, 0.1 mM EDTA, 1 mM DTT, 5% Glycerol, 0.5 mM PMSF, 10 µg/ML Aprotinin, 10 µg/mL Leupeptin, 0.1% NP-40, 1x HALT) was added to cell pellets. Cell pellets incubated on ice for 5 minutes before being centrifuged at 15,600 x g at 4°C for 10 minutes. The supernatants were transferred to a fresh 0.5 mL tube as cytoplasmic fractions. The cell pellets containing nuclei received 100 µL of High Salt Buffer (20mM HEPES, pH 7.9, 0.4M NaCl, 1mM EDTA, 1mM EGTA, 1mM DTT, 0.5 mM PMSF, 10 µg/mL Aprotinin, 10 µg/mL Leupeptin, 1xHALT) and incubated for 30 minutes on ice. All fractions were sonicated for three 5-second on pulses followed by 5-second off pulses at 25% amplitude. Then, all samples were centrifuged for 10 minutes at 18,213 x g at 4°C. All supernatants were transferred to fresh 0.5 mL Eppendorf tubes and stored at -80°C until immunoblotting. To confirm the purity of subcellular fractionation, 20 µg of protein resolved onto 4-12% acrylamide PAGE gels. After transferring, nitrocellulose membranes were probed with goat anti β-actin as a loading control, rat anti α-tubulin as a cytoplasmic marker, rabbit anti Histone H3, rabbit anti V5 to visualize TurboID-NES, and streptavidin- 680 to visualize biotinylation. Secondary antibodies included goat-anti-rat 800, donkey-anti-goat 680, and donkey-anti-rabbit 800. All primary and secondary antibodies incubated for 1 hour at room temperature, and were probed serially to ensure the specificity of the antibodies for their target.

### 6.8 Sample preparation for Mass Spectrometry studies

From each sample, 1 mg of lysate was set aside for streptavidin affinity purification, 50ug of protein were reserved as WC fractions, and the remaining protein was aliquoted and reserved for quality-control studies. Quality control studies were conducted prior to submitting samples for MS analysis to confirm the presence of biotinylated proteins via western blot, equal loading via Coomassie, as well as ensure the specificity of streptavidin-purified preparations for biotinylated proteins via silver stain (Pierce, Thermo fisher #24612) and western blot.

Slightly modified from previous publications the AP samples were processed as follows: ^13,22^ 1 mg of protein derived from transduced and untransduced BV2 and N2A lysates were affinity-purified onto 83μL of magnetic streptavidin beads (Thermo #88817). Briefly, to each 1.5 mL Eppendorf LoBind tube, 1 mL of RIPA buffer (150 mM NaCl, 50mM Tris, 1% Triton X-100, 0.5% sodium deoxycholate, 0.1% SDS, pH 7.5) was added to the beads on rotation for 2 minutes at room temperature (RT). Using a magnetic stand (PureProteome, Millipore #LSKMAGS08), the buffer was removed from the beads. To each tube-containing-beads, 500μL of fresh RIPA lysis buffer was added before adding 1mg of protein. The samples incubated at 4°C overnight on a rotator. Samples were then briefly centrifuged, placed on the magnetic stand, and the supernatants were preserved and frozen at -20°C. After incubation, the beads containing the biotinylated proteins underwent series of washing procedures at RT. Beads were washed twice with 1 mL of RIPA lysis buffer for 8 minutes, 1 mL 1M KCL for 8 minutes, rapidly (∼10 seconds) in 1 mL of 0.1 M sodium carbonate (Na_2_CO_3_), rapidly in 1 mL of 2M urea in 10mM Tris-HCl (pH 8.0), and twice with 1 mL RIPA lysis buffer for 8 minutes. 8-minute washing steps took place on a rotator, whereas rapid washing procedures took place on the magnetic stand using pipette rinsing to briefly mix the samples. Each buffer removal step was performed via manual pipetting, as aspirating the buffer with vacuum systems may deplete bead volume. The beads were resuspended in 1 mL of 1x PBS before being transferred to a new Eppendorf Lo Bind tube where they were washed once more with 1 mL of 1x PBS. Finally, AP samples were resuspended into 83μL of PBS, wherein 8μL of beads-containing solution was transferred to a new tube and placed on a magnetic stand for >2 minutes. The remaining beads-containing solution was preserved. The PBS was removed and replaced with 30μL of 4x Laemmli protein buffer (Bio-Rad #1610747) supplemented with β- mercaptoethanol, 2mM biotin, 20mM dithiothreitol (DTT). Beads then incubated at 95°C for 15 minutes and 20μL were reserved for Western Blot verification of biotinylated proteins via Streptavidin 680 and 10μL were reserved for separate silver stain to verify minimal nonspecific binding between untransduced AP samples and transduced AP samples.

To confirm the quality, equal loading, and biotinylation of proteins in the WC samples, 10μg of protein from each sample resolved onto a 4-12% polyacrylamide gel for Coomassie Blue staining, and 20 µg of protein was resolved on a separate gel to probe for actin and biotinylated proteins via immunoblotting. For Coomassie staining, gels were fixed in 50% methanol and 10% glacial acetic acid for 1 hour at RT in a sealed container on a rocking incubator. The gels were stained for 20 minutes (0.1% Coomassie Brilliant Blue R- 250, 50% methanol, 10% glacial acetic acid). Finally, a destaining solution (40% methanol, 10% glacial acetic acid) was applied at room temperature while gels incubated with gentle rocking. Coomassie gels were imaged in the 700nm channel on the LiCor Odyssey imaging system. Confirmation of biotinylated proteins and equal loading of WC samples took place by immunoblotting with streptavidin-conjugated fluorophore 680 (Strep680) and Gt anti actin, respectively.

### 6.9 Peptide Digestion & Cleanup

Sample preparation for MS was performed according to our laboratory protocols modified from previous publications^30,73–79^. Briefly, 50 µg of protein from each cell lysate sample was digested. Samples were reduced with 5 mM dithiothreitol (DTT) at room temperature for 30 minutes on a rotor, followed by alkylation with 10 mM iodoacetamide (IAA) at room temperature for 30 minutes on a rotor in dark. Samples were diluted with 50 mM of ammonium bicarbonate (ABC) to 4 M urea prior to undergoing overnight digestion with 2 µg of lysyl endopeptidase (Lys-C) (Wako #127-06621) at room temperature. Samples were then diluted to reduce the concentration of Urea to 1 M prior to trypsin digestion. Each sample received 2 µg of trypsin (Thermo, 90058) and were incubated overnight at room temperature. Acidifying buffer was added to the peptide solution for a final concentration of 1% formic acid (FA) and 0.1% trifluoroacetic acid (TFA) to stop the trypsin digestion. HLB columns were used to desalt samples (Waters #186003908). The samples were dried overnight using a centrifugal vacuum concentrator (SpeedVac Vacuum Concentrator).

AP samples underwent on-bead digestion. Beads were resuspended in 150 µL ABC. Application of DTT to a final concentration of 1 mM reduced the samples during a 30- minute room-temperature incubation on a rotator. A 5 mM application of IAA alkylated the samples during a 30-minute incubation in the dark on a rotator. To each sample, 0.5 μg of LysC was added before incubating overnight at room temperature on a rotator. The digestion was completed overnight at room temperature on a rotator with the addition of 1 µg of trypsin to the samples. After overnight digestion, the samples were treated with acidifying buffer to stop the trypsin-mediated digestion. HLB columns were used to desalt samples. The samples were dried using the SpeedVac.

### 6.10 Liquid Chromatography and Mass Spectrometry

All samples were analyzed on the Evosep One system using an in-house packed 15 cm x 150μm i.d. capillary column with 1.9 μm Reprosil-Pur C18 beads (Dr. Maisch, Ammerbuch, Germany) using the pre-programmed 44 min gradient (30 samples per day). Mass spectrometry was performed with a Q-Exactive Plus (Thermo) in positive ion mode using data-dependent acquisition with a top 20 method. Each cycle consisted of one full MS scan followed up to 20 MS/MS events. MS scans were collected at a resolution of 70,000, 400-1600 m/z range, 3x10^6 AGC, 100 ms maximum ion injection time. All higher energy collision-induced dissociation (HCD) MS/MS spectra were acquired at a resolution of 17,500 (1.6 m/z isolation width, 28% collision energy, 1×10^5 AGC target, 100 ms maximum ion time). Dynamic exclusion was set to exclude previously sequenced peaks for 30 seconds within a 10-ppm isolation window. MS raw files of WC and AP samples were searched together using the search engine Andromeda integrated into MaxQuant (Ver 1.6.17.0). Raw files were searched against the 2020 Uniprot *murine* database. Variable modifications include methionine oxidation, N-terminus acetylation, and deamidation of glutamine and asparagine residues. Fixed modifications include carbamidomethylation of cysteine residues. Only peptides with up to 2 missed cleavages were considered in the database search. Additional search parameters included a maximum peptide mass of 6000 Daltons (Da) and the minimum peptide length of 6 residues. Peptide spectral match (PSM) false discovery rates (FDR), protein FDR, and site FDR were all set at 1 percent.

### 6.11 Data normalization and filtering, Principal Component Analyses, Differential Expression Analyses, Clustering, Gene Set Enrichment Analysis

To analyze large datasets generated by MS, we used principal component analysis (PCA) as a dimension reduction strategy, differential expression analysis (DEA) was used to identify significant differences in protein intensities between samples, clustering analyses (k-means) identified discrete proteins with related abundance values within and across samples, and gene over-representation analysis functionally annotated enriched groups of proteins.

#### Filtering missingness, data normalization and log transformation

Label-free quantification intensities and raw intensity values were uploaded onto Perseus (Ver 1.6.15) for analyses. Categorical variables were removed, intensity values were log-2 transformed, transduced AP intensity values were normalized to TurboID intensity to adjust for variability in TurboID expression, and data were in general filtered based on 50% missingness across group of samples that were selected for each analysis. Missing values were imputed from normal distribution.

#### PCA

Principal component analyses of LFQ mass-spectrometry data were performed and visualized using SPSS (IBM Statistics, Version 28.0.1.0). The PCA on cytokine profiling with Luminex were performed using the PCA function from the Monte Carlo Reference-based Consensus Clustering (M3C) library (Bioconductor, Version 3.15).^80^

#### DEA

Differential Expression analysis (DEA) were performed in Perseus using students two sample t-test comparisons (unadjusted *p* value ≤0.05). For comparisons within AP samples, including untransduced AP vs. transduced AP and transduced N2A AP vs. transduced BV2 AP, and transduced BV2+LPS AP vs. transduced BV2 AP, TurboID-NES- normalized intensity values were used. For within WC comparisons, including transduced BV2+LPS WC vs transduced BV2 WCs, LFQ intensity values were used. Differentially enriched proteins were visualized as volcano plots with Prism (GraphPad, Ver 9.3.1 for Windows, San Diego, California USA, https://www.graphpad.com). The DEA of cluster-level significant changes in response to LPS (**SF 5**) was performed using the average intensity values of all proteins within a specific cluster for each biological replicate within a group. Then, the average cluster-intensity across replicates within an experimental group was taken. The average cluster-level responses to LPS were compared between whole cell and transduced AP groups with Log2FC DEA using an unadjusted p value ≤ 0.05, visualized as asterisks.

#### Clustering

To identify discrete groups of differentially enriched proteins across samples or associated with LPS treatment, we performed K-means clustering. The elbow method was used to determine the optimal number of clusters for K-means clustering by using Integrated Differential Expression and Pathway analysis (iDEP Ver .95 (http://bioinformatics.sdstate.edu/idep). Clustering analyses were visualized as heatmaps generated using Morpheus (Broad Institute, Morpheus, https://software.broadinstitute.org/morpheus).

#### Gene set enrichment analysis

Gene set enrichment analysis (GSEA) of differentially expressed proteins was performed using over-representation analysis (ORA) with the software AltAnalyze (Ver 2.0). Fisher exact significance threshold of p value ≤ 0.05 (Z-score greater than 1.96) was used to identify significant gene ontologies. Functional annotation of proteins biotinylated by TurboID-NES which significantly differ by cell-type (Fig 4C, 4E) took place by using enriched N2A and BV2 proteins (404 and 936 proteins, respectively) as input lists and the list of proteins identified in the AP dataset as the background (2277 proteins). Over-represented terms and their corresponding Z-scores were visualized as bar graphs using Prism. SynGO was used to identify unique synaptic terms in N2A-enriched proteins (Fig 4D)^81^. To functionally annotate K-means clusters (Fig 2B, Fig 5A), lists of gene symbols associated with each K-means cluster were input into AltAnalyze for ORA. The resulting z-scores underwent gene GO and KEGG hierarchical cosine-cosine clustering, and each cluster was functionally annotated and the associated z-scores were represented in heatmaps. Mapping gene-associated risk loci with Alzheimer’s disease, Parkinson’s disease and Amyotrophic Lateral Sclerosis relevant terms (**SF 3**), MAGMA dataset sheets were directly obtained from a previous review publication from our lab.^68^

### 6.12 Mitochondrial oxygen consumption in living BV2 cells

To determine if TurboID-NES expression, LPS treatment, or biotin supplementation impacts mitochondrial function in BV2 cells, we directly measured oxygen consumption rates (OCAR) and extracellular acidification (ECAR) parameters using a mitochondrial stress test in a Seahorse XFe96 extracellular flux analyzer (Agilent). In a 96-well cell microplate, 5,000 transduced or untransduced BV2 cells were seeded in 80 µL of complimented growth media and incubated at room temperature in sterile cell culture hood for 1 hour. After cells adhere to the bottom of the well, 120 µL of complimented growth media was added and cells incubated overnight at in the cell culture incubator at 37° C and 5% CO_2_. After 24 hours incubation, cells underwent a complete media change and were exposed to 1000ng/mL LPS or PBS as a vehicle control for 48 hours. The sensor cartridge was hydrated overnight at 37° C and 0% CO_2_ using 200 µL of sterile deionized water added to each well of the utility plate. After incubating overnight without CO_2_, water was removed from the utility plate and 200 µL of calibrant pre-warmed to 37°C was added to each well. The sensor plate incubated in calibrant at 37° in a CO_2_-free incubator for 1 hour prior to loading the cartridge. Prior to assay, cells were washed thrice with 180 µL of Seahorse Media (Phenol-free 5mM HEPES Seahorse XF media, 10mM glucose, 2mM L-glutamine, and 1 mM sodium pyruvate, pH 7.4). Cells incubated in a CO_2_-free incubator for 1 hour at 37° C. Sensor cartridges were loaded with 20 µL of Oligomycin (1.5 µM / well), 22 µL of carbonyl cyanide-4 (trifluoromethoxy) phenylhydrazone (FCCP) (0.75 µM / well), and 25 µL of Rotenone and Antimycin (0.5 µM / well) and calibrated. After calibration, the seahorse-assay plate containing BV2 cells was run using the mitochondrial stress test in Wave (Version 2.6). Cells were then stained with Hoechst 33342 dye for 20 minutes and imaged using ImageXpressTM Micro Confocal imaging, and cells were counted using the Find Blobs feature of MetaExpress Software’s Count Nuclei Application. Oxygen Consumption Rates (OCR) and Extracellular Acidification Rates (ECAR) were normalized to cell counts.

### 6.13 Cytokine profiling of supernatants

Cytokine profiling was performed as previously published with modifications.^22^ Luminex multiplexed immunoassays (Cat # MCYTMAG-70K-PX32) quantified cytokines from cultured supernatants of transduced and untransduced BV2 and N2A cells receiving LPS or PBS SHAM stimulus (n = 6 / group). The cytokine panel detected Eotaxin, GM-CSF, INF-γ, IL-1a, IL-1b, IL-2, IL-4, IL-3, IL-5, IL-6, IL-7, IL-9, IL-10, IL-12p40, IL12p70, LIF,IL-13, LIX, IL-15, IL-17, IP-10, KC, MCP-1, MIP-1a, MIP-1b, M-CSF, MIP-2, MIG, RANTES, VEGF, and TNF-α. The average background intensity reading from each cytokine panel was subtracted from the raw cytokine abundance values and negative values were imputed to zero. To find appropriate loading volume for samples, linear ranging was performed as previously published^82^ and 24% of total sample volume was loaded. Assays were read on a MAGPIX instrument (Luminex).

## Supporting information

Supplemental Datasheet 1

Supplemental Datasheet 2

## Author Contributions

**Conception and Design:** Sydney Sunna, Sruti Rayaprolu, Srikant Rangaraju, Nicholas T. Seyfried. **Data Collection:** Sydney Sunna, Christine Bowen, Hollis Zeng**. Contribution of data and analytic tools:** Duc M. Duong, Pritha Bagchi, Qi Guo, Prateek Kumar, Christine Bowen, Sara Bitarafan, Levi Wood. **Analysis:** Sydney Sunna, Aditya Natu, Christine Bowen, Srikant Rangaraju. **Writing:** Sydney Sunna, Christine Bowen, Srikant Rangaraju. **Editing:** Sydney Sunna, Christine Bowen, Sruti Rayaprolu, Prateek Kumar, Pritha Bagchi, Sara Bitarafan, Aditya Natu, Levi Wood, Nicholas T. Seyfried, Srikant Rangaraju.

## Acknowledgements

Research in this publication is supported by the National institute of Aging of the National Institutes of Health: F31AG071319 (S. Sunna), F31AG074665-01 (C. Bowen), T32GM135060-03 (C. Bowen), F32AG064862 (S. Rayaprolu), R01AG075820 (S. Rangaraju), RF1AG071587 (S. Rangaraju), R01NS114130 (S. Rangaraju), R01AG061800 (N.T.S), U01AG061357 (N.T.S). This research project was supported in part by the Viral Vector Core of the Emory Center for Neurodegenerative Disease Core Facilities. This study was also supported in part by the Emory Flow Cytometry Core (EFCC), one of the Emory Integrated Core Facilities (EICF), and is subsidized by the Emory University School of Medicine. Additional support was provided by the National Center for Georgia Clinical & Translational Science Alliance of the National Institutes of Health under Award Number UL1TR002378. The content is solely the responsibility of the authors and does not necessarily reflect the official views of the National Institutes of Health. We would like to acknowledge Dr Eric Dammer (Emory University) for helping us with data analyses.

## Availability of data

Data are available via ProteomeXchange with dataset identifier PXD036744. Processed data are available as supplemental data files.^83^

## Supplemental

**SF1.**
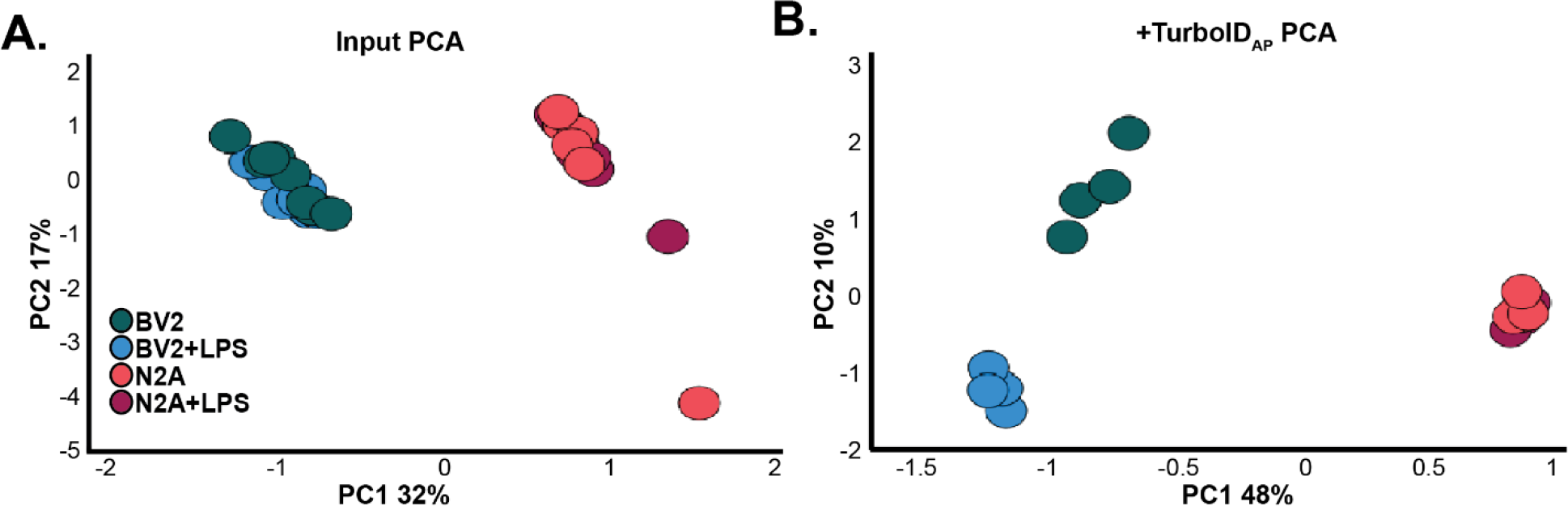
Dimension reduction of whole-cell and transduced AP samples. **A.** Principal component analysis (PCA) of whole-cell samples. Principal component 1 (PC1) accounts for ∼32% of variance across which samples distinguish by cell type. **B.**PCA on TurboID labeled and biotin enriched samples indicates that PC1 accounts for ∼48% of the variance and along this axis samples cluster by cell-type. PC2 accounts for 10% of the variance, along this axis, BV2 samples and not N2A samples cluster according to LPS treatment.

**SF2.**
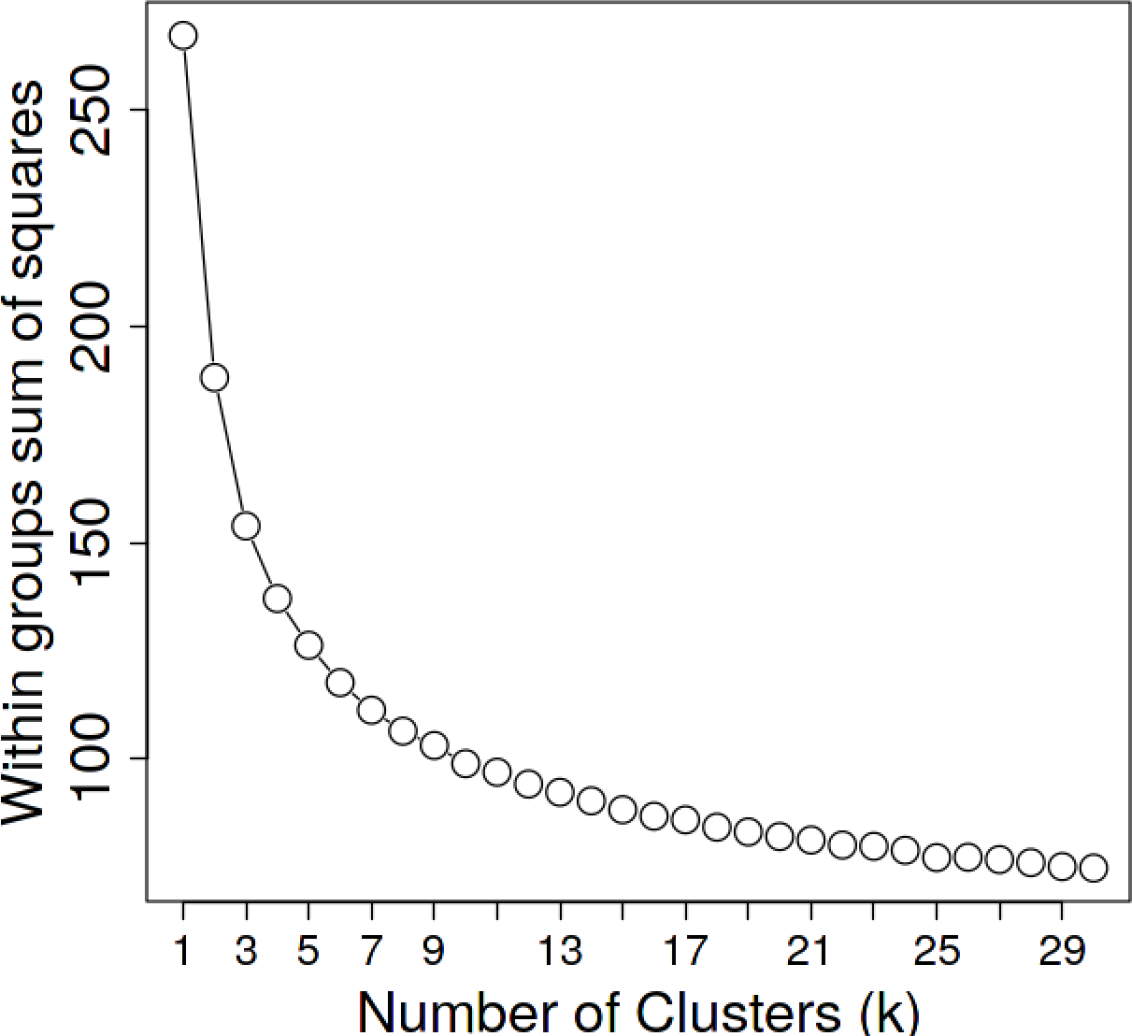
Optimization of cluster number using the Elbow-Curve Method. Mathematical determination of the optimal number of clusters for the clustering analysis performed in Figure 3.

**SF3.**
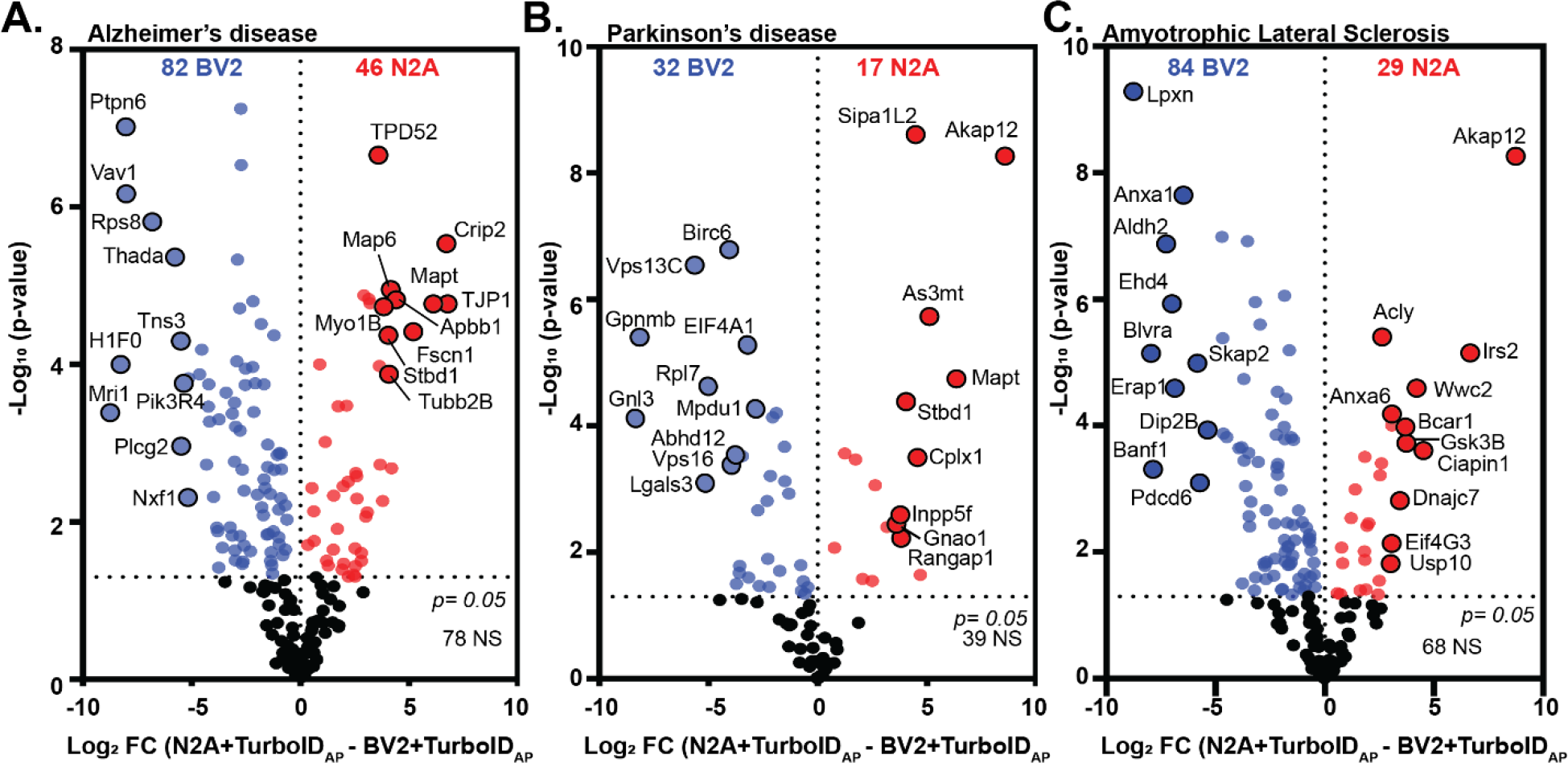
TurboID labels cellularly distinct proteins with relevance to neurodegenerative disease. Volcano plot representations of cellularly-distinct proteins labeled by TurboID (derived from significant DEP dataset in Fig 4A), and respective disease relevance. **A.** Mapping proteins onto the Alzheimer’s disease (AD) AD MAGMA dataset, there are 82 AD-associated proteins enriched in biotin-labeled BV2 proteins and 46 AD-related proteins enriched in biotin-labeled N2A proteins. **B.** Mapping biotin-labeled and cellularly distinct proteins onto the Parkinson’s disease (PD) MAGMA dataset, there are 32 PD-relevant proteins in the BV2 biotin-labeled proteome and 17 PD-relevant proteins in the N2A biotin-labeled proteome. C. Mapping proteins onto the Amyotrophic lateral Sclerosis (ALS) and Frontotemporal dementia (FTD) MAGMA dataset, there are 84 ALS/FTD-relevant BV2 proteins and 29 proteins in the N2A biotin-labeled proteins

**SF4.**
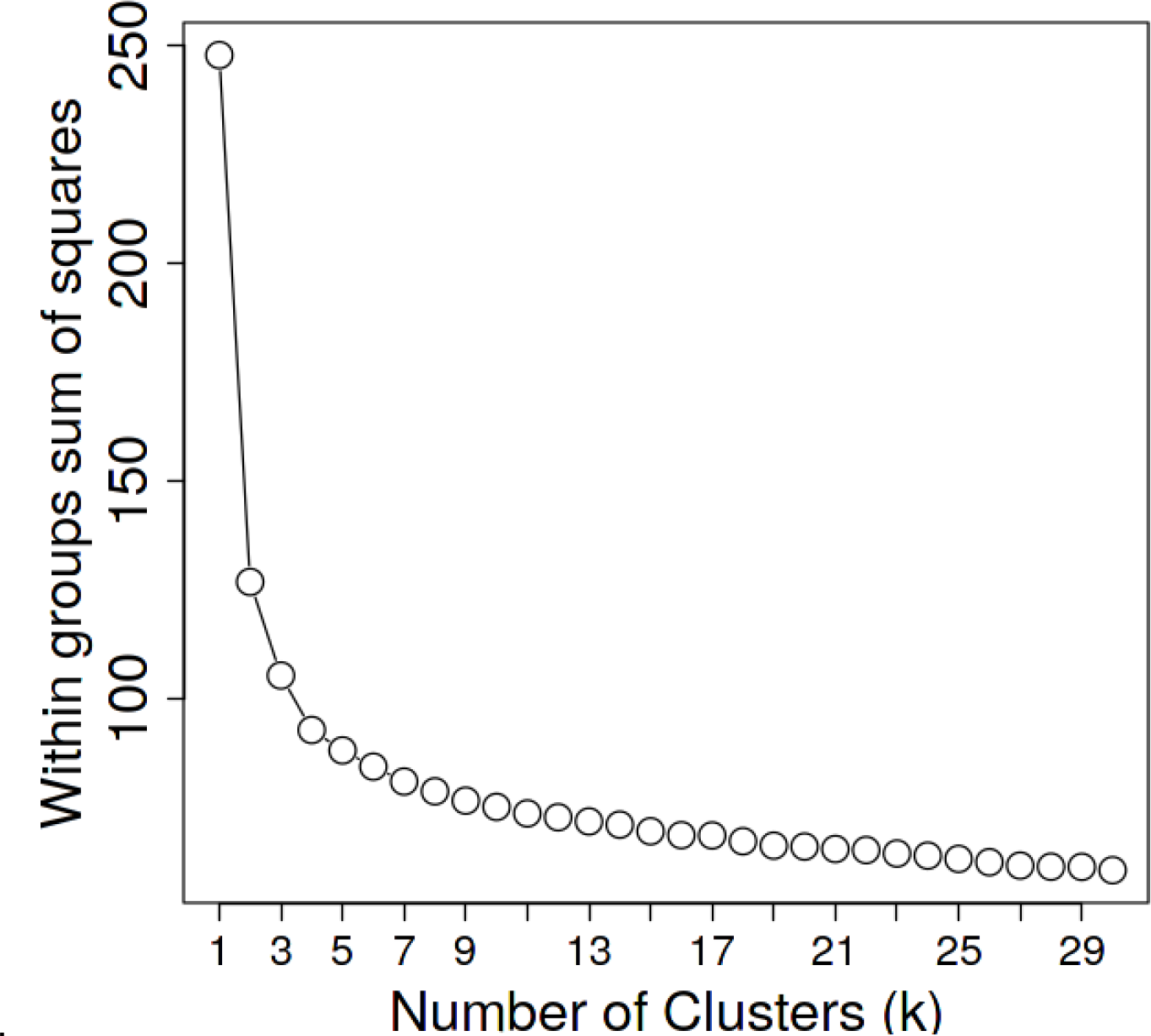
Optimization of cluster number using the Elbow-Curve Method. Mathematical determination of the optimal number of clusters for the clustering analysis performed in Figure 6

**SF5.**
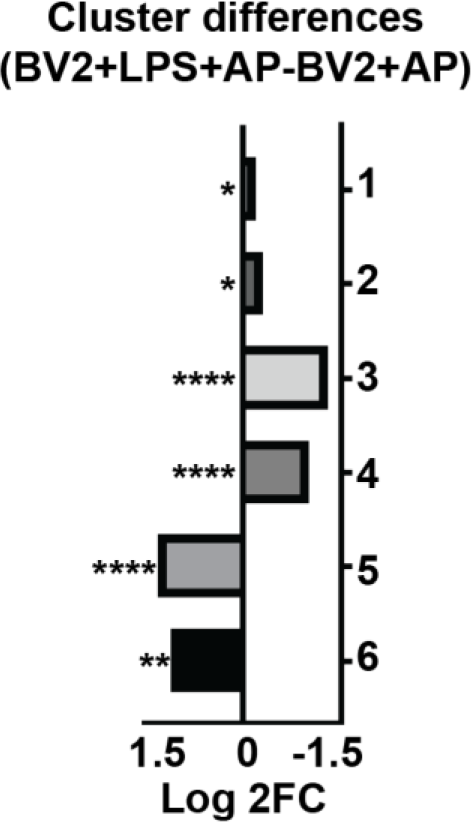
Log 2 FC of LPS driven changes between AP samples across all clusters. Bar-graph representation of the average Log2FC between BV2+TurboID+LPS AP and BV2+TurboID AP samples at the cluster-level. Clusters 1 and 2 show a minimal impact of LPS on the transduced AP proteomes (***Cluster 1:*** Log2FC = -0.2, p = 0.015 **& *Cluster 2:*** Log2FC = -0.3, p = 0.015). Clusters 3 and 4 show a robust LPS-mediated decrease in proteins in BV2 AP (***Cluster 3:*** Log2FC = -1.3, p = 1.49 E^-5^ **& *Cluster 4:*** Log2FC = -1.0, p = 9.55 E^-5^). Clusters 5 and 6 identify an LPS-mediated increase in protein abundance within BV2 AP samples (***Cluster 5:*** Log2FC = 1.3, p = 1.37 E^-5^ **& *Cluster 6:*** Log2FC = 1.1, p = 0.005).

**SF6.**
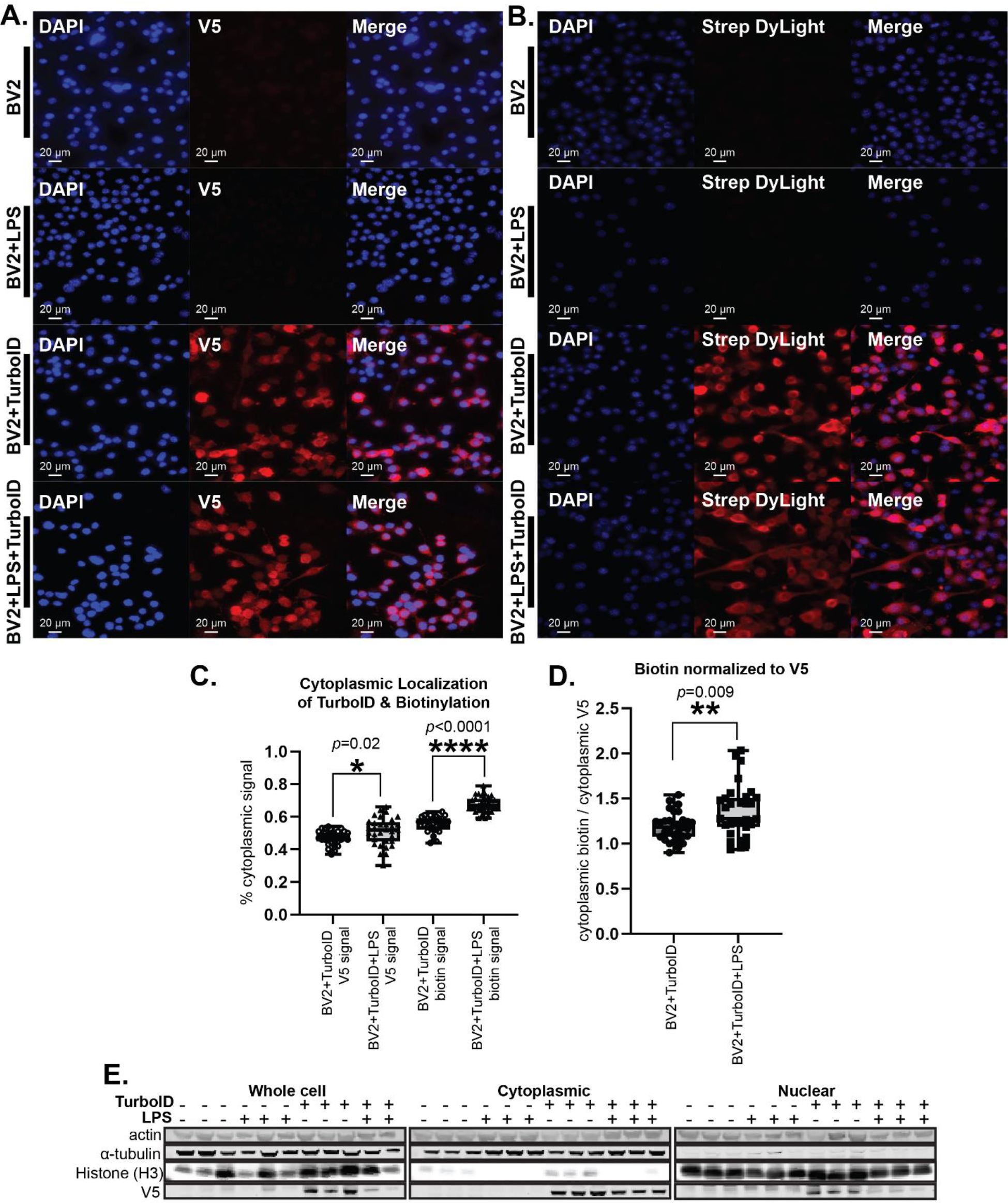
LPS challenge slightly increases cytosolic direction of TurboID and biotinylation of proteins. **A.** Immunofluorescence (IF) visualizing TurboID (V5, red) and nuclei (DAPI, blue) localization in transduced and untransduced BV2 cells with and without LPS challenge. Localization of V5-TurboID remains cytosolic under LPS challenge. **B.** IF visualizing biotinylation of proteins (StrepDylight, red) and nuclei (DAPI, blue) in contexts of LPS challenge in transduced and untransduced BV2 cells. Biotinylation of proteins remains cytosolic under LPS challenge. **C.** Colocalization analysis of the area of cytoplasmic V5 signal (*left two* box-*plots*) and cytoplasmic biotinylation signal (*right two box-plots*) indicates a significant increase in cytosolic V5 and biotinylated proteins with LPS challenge. Significant p values were determined using the two-tailed Mann-Whitney test. **D.** Biotinylation signals normalized to V5 signals within the cytoplasm. After normalizing the biotinylation intensity values to V5 intensity values, there is a significant increase in cytosolic biotinylation of proteins with LPS. Significant p values were determined using the two-tailed Mann-Whitney test. **E.** Western blot (WB) verification of subcellular-fractionation experiments of transduced and untransduced BV2 cells with and without LPS challenge (n=3 /group). Using β-actin as a loading control, α-tubulin as a cytoplasmic marker, histone H3 as a nuclear marker, and V5 as a marker for TurboID, we can confirm purification of sub-cellular fractions with the decrease in α-tubulin signal in the nuclear fraction as compared with the whole cell and cytoplasmic fractions and a decrease in histone H3 in the cytoplasmic fraction as compared with the whole cell and cytoplasmic fractions. With actin signal intensity remaining constant as a loading control, there is no apparent difference in cytoplasmic V5 intensity with LPS challenge.

